# Integrative analysis of pooled CRISPR screens provide functional insights into AD GWAS risk genes

**DOI:** 10.1101/2025.07.17.660041

**Authors:** Rebecca Y. Wang, Glenn Wozniak, Xiaoting Wang, Meer Mustafa, Malak El Khatib, Elias Kahn, Peter Heutink, Hua Long, Sara Kenkare-Mitra, Arnon Rosenthal, Zia Khan, Julia A. Kuhn, Daniel R. Gulbranson

## Abstract

Emerging research has implicated Alzheimer’s disease (AD) pathology with dysregulation of many key pathways in microglia, including lipid transport and metabolism, phagocytosis of plaques, and lysosomal function. However, the exact mechanisms underlying these pathways remain poorly understood. Leveraging high-throughput CRISPR screens to understand the interplay between these pathways may enable novel therapeutic strategies for AD and other neurological diseases. Here, we constructed activation and interference CRISPRa/i libraries targeting 203 genes, 71 of which were identified through neurodegenerative GWAS, and 132 additional genes linked to microglial functions. We used this library to conduct pooled CRISPRa/i screens across a range of functional assays relating to lipid metabolism and lysosomal function using a monocytic cell line, THP-1. We identified a core set of lipid and lysosome mediators and validated a subset in primary macrophages. To gain insights into transcriptional states modulated by these genes we also applied the CRISPRa/i libraries to Perturb-seq, enabling us to capture transcriptomic changes. Through non-negative matrix factorization, we identified five gene programs altered by our perturbation library. We then used an integrative analysis of functional screen data with Perturb-seq data that enabled us to uncover novel functions and genetic relationships between perturbations. This multidimensional resource links genetic perturbations to phenotypes and transcriptional programs, establishing a scalable framework for systematic gene discovery in neurodegeneration and beyond.

Graphical Abstract

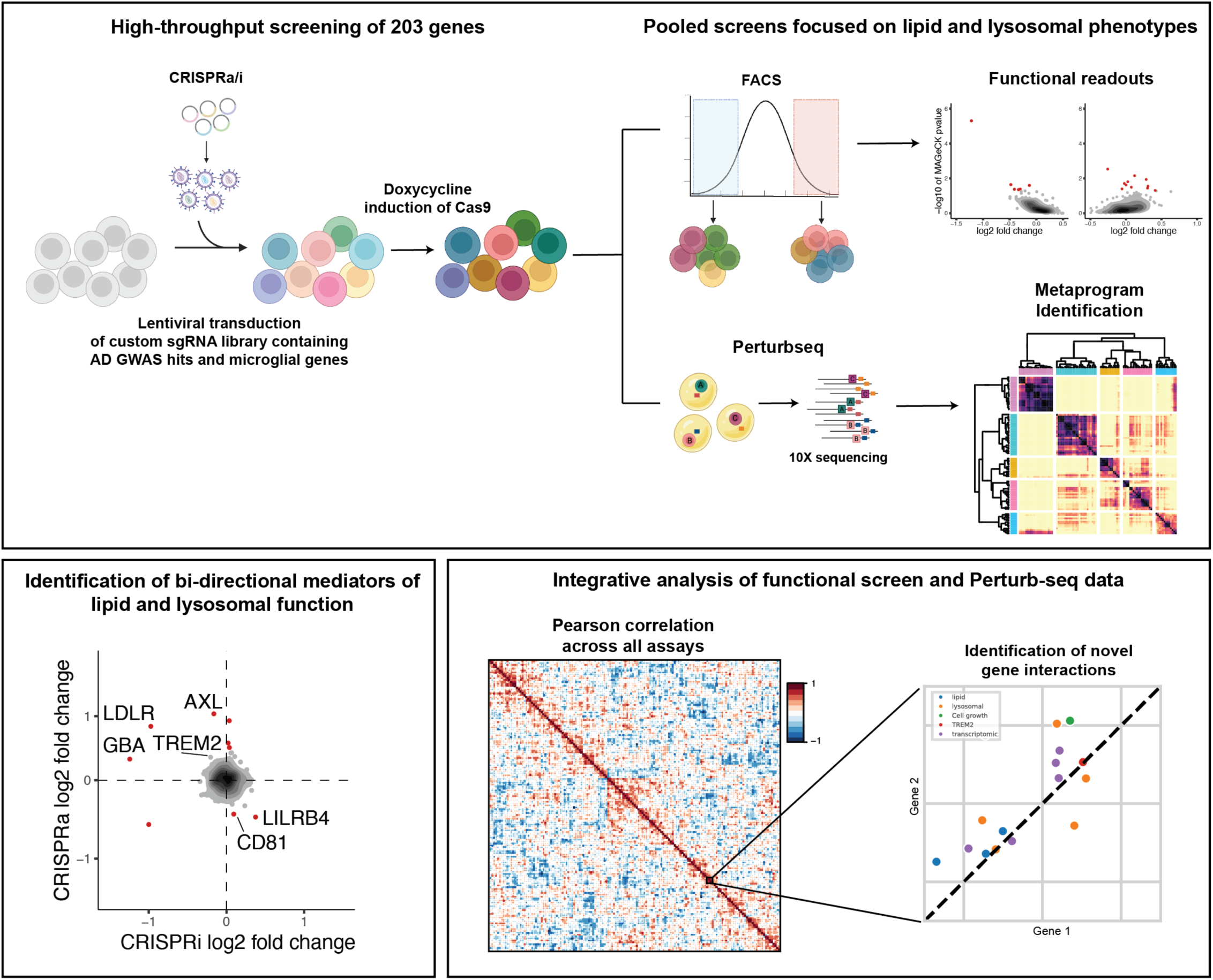

## Introduction

Human genetic studies have implicated microglial genes with neurodegenerative diseases, including Alzheimer’s disease (AD), suggesting that dysfunctional microglia may be a key contributor to disease development or progression^1–3^. Microglia, the brain’s resident immune cells, derived from the myeloid lineage, play a vital role in supporting brain health in part by responding to and mitigating the impact of injuries to the brain^4–6^. In AD, microglia migrate to lesions containing extracellular amyloid beta (Aβ) where they sequester, phagocytose and degrade Aβ^7,8^. However, most approved or investigational therapeutics for AD target the pathology present in the disease rather than modulating the underlying cause^9^, which in some cases is impaired microglia function^10–12^. One promising target for modifying microglia function is TREM2, a master microglia regulator that functions as a lipid receptor^13,14^. TREM2 is activated by ligand binding, triggering a downstream signaling cascade that results in increased microglia survival, proliferation, phagocytosis, and lysosomal degradation^15–18^. While direct TREM2 activation is currently tested in clinical trials for neurodegenerative diseases^19^, finding new upstream and downstream regulators of TREM2 signaling would identify alternative target candidates for the treatment of neurodegenerative diseases. Identifying additional targets capable of modifying microglia function could advance AD drug development.

In AD, one critical function of microglia is the clearance of Aβ^4,7^, which occurs in the lysosome after phagocytosis^20–22^. Some studies have shown that a build-up of Aβ is a consequence of lysosomal dysfunction^23–25^. Consistent with this idea, loci in or near multiple lysosomal genes including *TMEM106B*, *GRN*, *ABCB9*, *BLOC13S*, *CYB561*, *IDUA*, *SNX1*, and *WDR1* are associated with AD^1^. Enhancing lysosomal function may offer therapeutic benefits in AD or other neurodegenerative diseases. In addition to its role in protein degradation after phagocytosis or during autophagy, the lysosome plays an important role in maintaining lipid homeostasis^26,27^. Lipoproteins bind to cell surface receptors and are internalized into the endosome, which then fuses with the lysosome^28^. Once in the lysosome, the low pH facilitates the dissociation of the lipoprotein from its receptor, which then recycles back to the cell surface, while the lipoprotein is degraded into its constituent parts, mostly apolipoproteins, triglycerides, cholesterol, and phospholipids^28^. Notably, one apolipoprotein, ApoE, is the strongest genetic risk factor for AD, and other lipid-related genes such as *ABCA7*, *CLU*, and *PLD3* are also implicated in AD by human genetic studies, suggesting that lipid homeostasis may be a key contributor to AD^1^. Interestingly, a recent genome-wide screen identified lipid homeostasis pathways as top regulators of endogenous protein aggregation, suggesting that lipid homeostasis and lysosomal degradation pathways may converge to drive neurodegenerative diseases^29^.

Although many loci identified in genome-wide association studies (GWAS) are linked to microglia genes, some with known roles in lipid homeostasis or lysosomal function, two critical pathways for microglia biology, many others remain poorly understood. In order to better understand the role of these genes in key microglia processes, we designed custom activation and interference sgRNA libraries based on human genetic links to neurodegenerative diseases. We used these sgRNA libraries in a range of functional assays relating to TREM2 expression, lipid uptake and metabolism, and lysosomal function. Notably, we identified TGFBR2 as strong regulator of surface TREM2 levels, which is consistent with previous GWAS that implicated this loci as being associated with CSF TREM2 levels in humans^30^. We also identified a core set of genes, including TREM2, that regulate lipid homeostasis and lysosomal function and validated a subset of these genes in primary human macrophages (hMΦ).

Because single-cell RNA sequencing studies have begun to highlight the existence of different microglia subtypes present in disease, a phenotype that is primarily defined by gene expression, we explored the use of Perturb-seq as a platform for performing screens with transcriptomic readouts. In total, we performed 24 CRISPRa/i screens, generating a rich multimodal dataset for integrative analysis. With the combination of both functional and transcriptomic readouts, we capture a broader range of gene functions, allowing us to identify novel genetic relationships between perturbations. Through this analysis, we show that the combination of functional and transcriptomic readouts enables unbiased clustering of genes based on their behavior in the component screens, enabling us to predict functions of poorly characterized genes at scale.

## Results

### Targeted screen for regulators of surface TREM2

To better characterize microglia genes relevant for neurodegenerative disease, we designed custom sgRNA libraries against human genetics-implicated microglia genes, primarily those whose gene product localizes to the cell surface, making it druggable with an antibody. This filtering resulted in 71 disease-implicated genes (Figure 1A). We added genes encoding cell surface receptors and intracellular proteins important for microglia functions resulting in a total of 203 genes (5 sgRNAs each) with 100 non-targeting control sgRNAs (Figure 1B). Since we wanted a scalable myeloid model enabling us to perform dozens of screens quickly, we chose to use THP-1 cells for screening. We generated tet-inducible dCas9-KRAB or dCas9-VPH cells by transducing THP-1 cells with lentivirus and then added our custom sgRNA library (Figure 1C).

**Figure 1:**
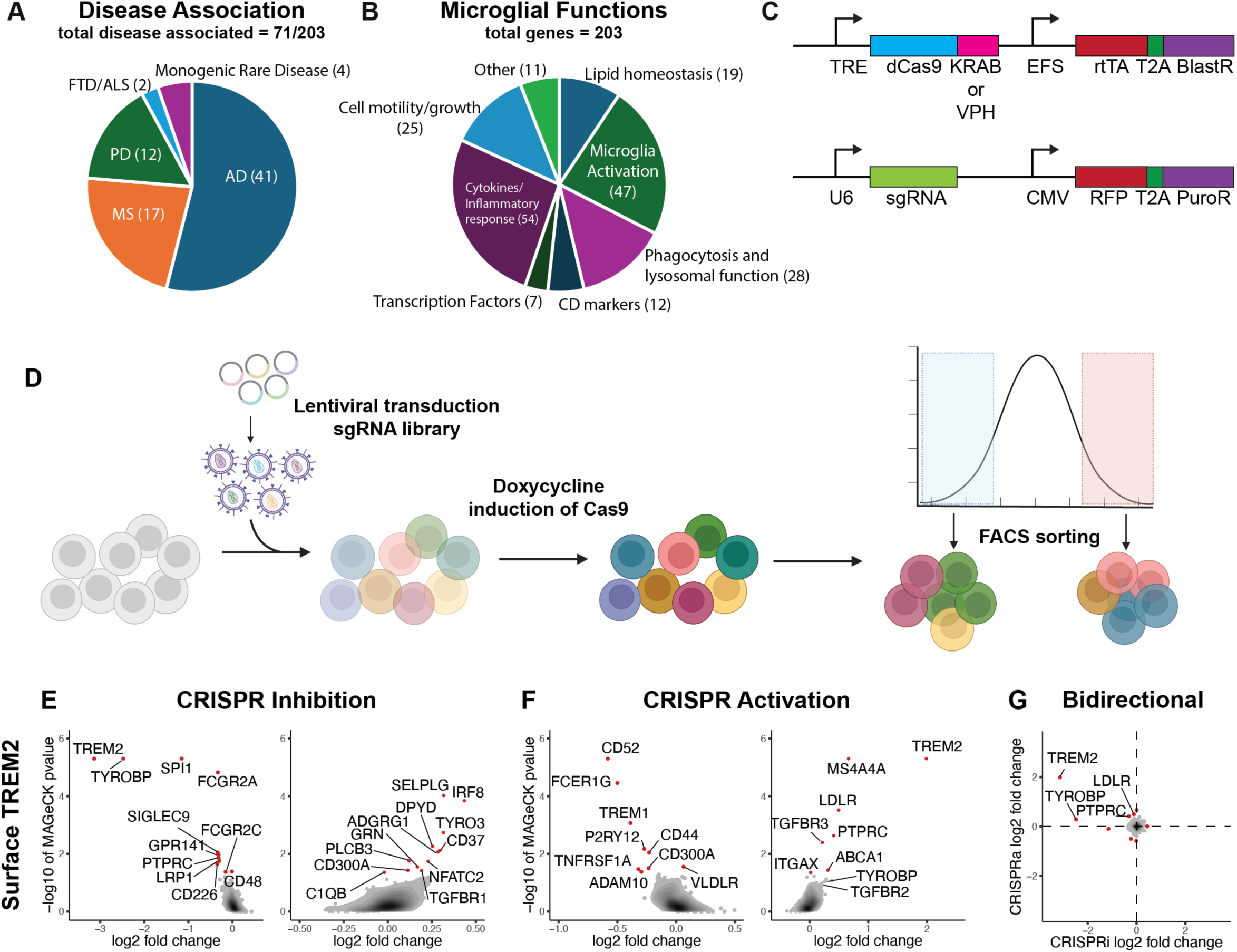
Paired CRISPRa/i screens identify regulators of surface TREM2 levels. A, B) Pie graphs showing the proportion of sgRNA targets in focused CRISPRa or CRISPRi sgRNA libraries that have been implicated by human genetic associations for neurodegenerative disease (A) or whose gene product has a given microglial function (B). C) Transgene design used to make stable CRISPRa or CRISPRi THP-1 cells (top) or to deliver sgRNA libraries (bottom). D) Schematic depicting the experimental workflow for performing surface TREM2 screening. E, F) Dot plot with an overlayed density plot of genes from CRISPRi (E) or CRISPRa (F) screens for surface TREM2 levels. The normalized log2 fold change in readcount of the sgRNA for a given gene with the median change in abundance in the high surface TREM2 population relative to the low surface TREM2 population is plotted on the x-axis against the -log10 P-value calculated by MAGeCK for negative enrichment (left) or positive enrichment (right) on the y-axis. Genes with a P-value >0.05 are labeled red. G) Log2 fold change as defined above for CRISPRi screen (x-axis) plotted against the log2 fold change for in the CRISPRa screen (y-axis).

To evaluate this model as a screening paradigm we induced dCas9 expression by growing the cells for 7 days in the presence of doxycycline, stained them with an antibody against TREM2, a critical mediator of myeloid cell function, and used fluorescence activated cell sorting (FACS) to collect the 10% of cells with either the highest or lowest surface TREM2 levels (Figure 1D). Frequencies of sgRNAs in each group were determined by next-generation sequencing (Supplementary Table 1). MAGeCK^31^ was used to calculate enrichment and significance of sgRNAs in one population relative to the other (Supplementary Table 2). As expected, reducing TREM2 with CRISPRi led to a robust decrease in surface TREM2 levels, whereas increasing TREM2 with CRISPRa led to robust increase in surface TREM2 levels in our screens (Figure 1E, F). Beyond *TREM2* itself, *MS4A4A* was the strongest regulator of surface TREM2 levels in our CRISPRa screen indicating a functional relationship between genes in the MS4A locus and TREM2. Consistent with this *in vivo* human data, our surface TREM2 CRISPRa screen identified both *TGFBR2*, as well as it’s coreceptor *TGFBR3* (Figure 1F). To validate this result, we treated THP-1 cells with TGF-β, a ligand for these receptors. This resulted in an increase in surface TREM2 levels, as well as increased soluble TREM2 levels in the media of these cultures (Figure S1A-C).

We reasoned genes whose knockdown (KD) or overexpression (OE) resulted in opposite phenotypes, which we refer to as bidirectional regulators, may be critical mediators of surface TREM2. Indeed, decreasing *TYROBP* expression, which encodes the TREM2 signaling effector DAP12, led to reduced surface TREM2 levels in our CRISPRi screen, whereas OE of DAP12 led to increased TREM2 levels (Figure 1E–G). *PTPRC*, which encodes CD45, a member of the protein tyrosine phosphatase family that is often used as a microglia marker, also bidirectionally regulated TREM2 in our screen (Figure 1G). LDLR, which encodes low-density lipoprotein receptor, a critical mediator of cellular cholesterol levels also bidirectionally regulated surface TREM2 in these screens (Figure 1G).

### Targeted screens for regulators of lipid homeostasis and lysosomal function

To better understand which genes in our sgRNA libraries regulate lipid uptake, we performed CRISPRa/i screens by treating THP-1 cells with pHrodo-labeled LDL and FACS sorting the cells with the highest or lowest fluorescent levels. *LDLR* and *SCARB1*, which are known regulators of LDL uptake^28,32^, were top hits in both the CRISPRa and CRISPRi screens, as well as the cholesterol efflux regulator *ABCA1* (Figure 2A–C). We also identified unexpected bidirectional regulators of LDL uptake including *TREM2*, *TGFBR3*, and *SQSTM1*, an autophagy regulator^33^. *LDLR* was a top bidirectional regulator of surface TREM2 (Figure 1E–G), and *TREM2* was a top bidirectional regulator of LDL uptake (Figure 2A–C), suggesting a feedback loop may exist in which LDLR regulates surface TREM2 levels, which in turn regulates LDL uptake. To gain insights into how specific the lipid uptake regulators were for LDL, we performed additional CRISPRa/i screens for pHrodo-labeled ApoE4 uptake. As in the LDL uptake screens, *LDLR*, *ABCA1*, and *SQSTM1* were top bidirectional regulators of ApoE4 uptake (Figure 2A–F). *SCARB1* and *TREM2* were specific regulators of LDL uptake, whereas *VLDLR* and *SIRPA* were specific regulators of ApoE uptake (Figure 2A–F). Not surprisingly, these screens also identified genes whose phenotype only appears in either the CRISPRa or CRISPRi screen. For example, upregulation by CRISPRa of *AXL*, which encodes a phagocytosis regulator^34^, led to increased LDL and ApoE4 uptake (Figure 2B, E). Similarly, upregulation of *CD68*, which encodes a lysosomal membrane protein^35^ led to more LDL and ApoE4 uptake in our screens, whereas reduction of *CD68* by CRISPRi only diminished ApoE4 uptake (Figure 2B, D–F). To clarify if modulation of these lipid uptake regulators altered steady-state lipid levels we performed CRISPRa/i screens using Bodipy, which stains neutral lipids. As expected, overexpression by CRISPRa of *ABCA1*, a lipid efflux transporter, diminished lipid levels validating the screen (Figure S2B). However, most of the genes identified in the lipid uptake screens were not hits in the Bodipy screen (Figure S2A–C), suggesting that genetic manipulation of these genes alters lipid uptake but not steady-state lipid levels.

**Figure 2:**
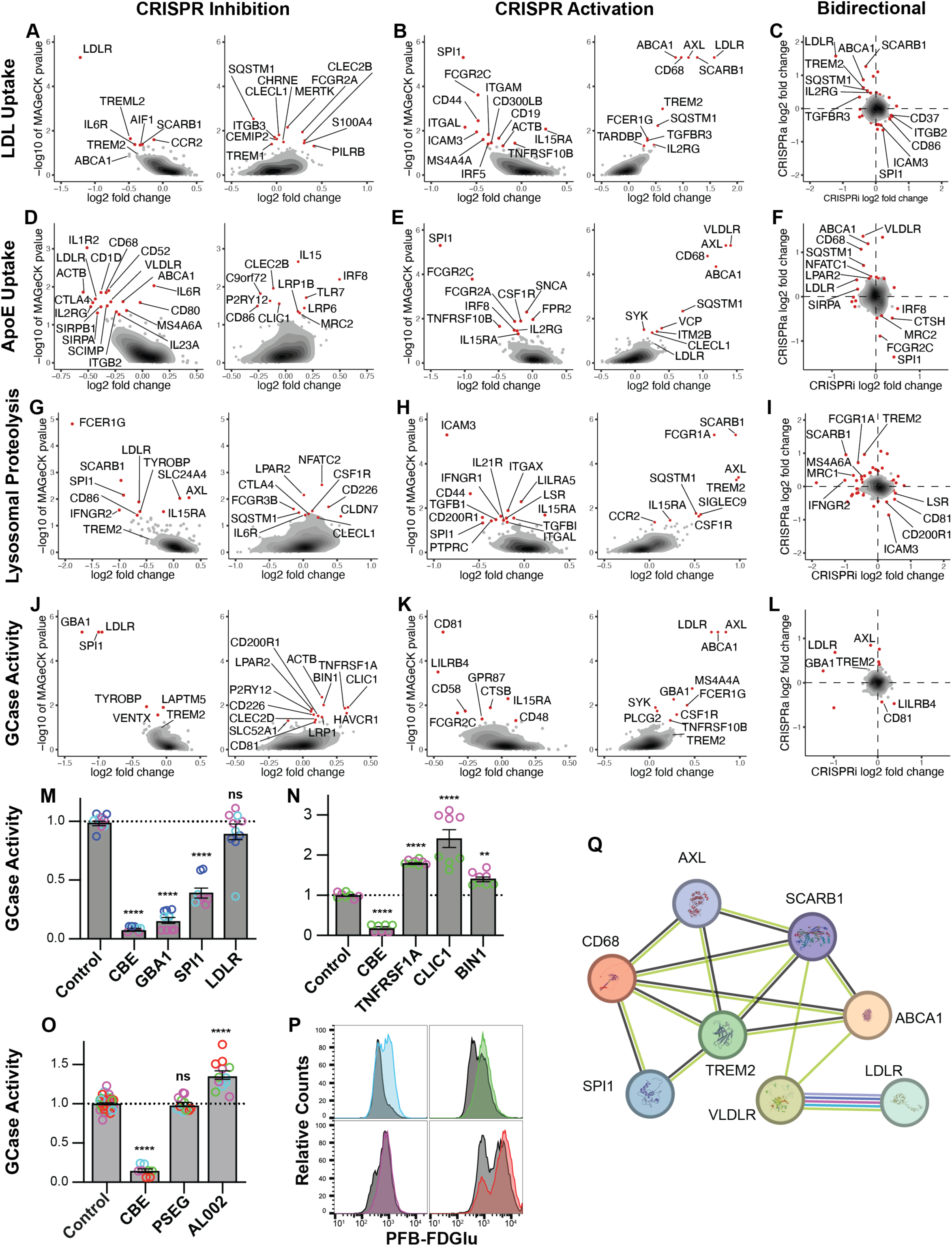
Paired CRISPRa/i screens identify regulators of lipid uptake and lysosomal function. A, B, D, E, G, H, J, K) Dot plot with an overlayed density plot of genes from CRISPRi (A, D, G, J) or CRISPRa (B, E, H, K) screens for LDL uptake (A, B), ApoE4 uptake (D, E), lysosomal proteolysis (G, H), or GCase activity (J, K). The normalized log2 fold change in readcount of the sgRNA for a given gene with the median change in abundance in the high fluorescent population relative to the low fluorescent population is on the x-axis. The -log10 P-value calculated by MAGeCK for negative enrichment (left) or positive enrichment (right) is on the y-axis. Genes with a P-value >0.05 are labeled red. C, F, I, L) Log2 fold change as defined above for CRISPRi screens (x-axis) plotted against the log2 fold change in the CRISPRa screens (y-axis). M–P) Primary human macrophages were electroporated with Cas9 protein and sgRNAs (M, N) or treated with an antibody (O, P) and GCase activity was measured by flow cytometry 1 h after treatment with PFB-FDGlu. The mean fluorescent signal from a well was dived by the mean fluorescent signal from all control wells. Dots indicate different wells, colors indicate different donors, error bars are the standard deviation; significance was calculated using one-way ANOVA with Holm-Sidak correction for multiple testing. **** P < 0.0001, ** P < 0.01, ns = not significant P) Histogram from cells of the well with the median fluorescent signal treated with a control antibody (PSEG) in gray or AL002, which are colored based on the donor they are derived from. Q) A String diagram generated by entering a curated list of genes from the lipid and lysosome screens using version 12.0 of string-db.org. Green lines connecting the proteins indicate the two proteins are frequently mentioned together in automated, unsupervised textmining. Black lines indicate correlated expression of the two genes across many experiments. Additionally, VLDLR and LDLR had known interactions from curated databases, experimentally determined interactions, protein homology, and were part of gene families whose occurrence patterns across genomes show similarities.

After LDL or ApoE4 is taken up by the cell, they are trafficked to the lysosome where they are degraded, releasing cholesterol^28^. To identify genes in our sgRNA libraries that regulate lysosomal function, we performed CRISPRa/i screens using a DQ-BSA assay. In this assay, proteases in the lysosome degrade BSA resulting in de-quenching of conjugated self-quenched fluorophores and increased fluorescence. Like in the previous screens, the top and bottom 10% of cells were FACS sorted based on their fluorescent signal and enrichment analysis of sgRNAs in the two populations was performed. Interestingly, *TREM2* and *SCARB1*, which were both top bidirectional regulators of LDL uptake, also bidirectionally regulated lysosomal proteolysis (Figure 2A– I). Similarly, OE by CRISPRa of *SQSTM1* or *AXL*, which led to increased LDL and ApoE uptake also led to higher lysosomal proteolysis (Figure 2A–I).

In addition to proteins, the lysosome degrades lipids such as ceramides. GCase, which is encoded by *GBA1*, is a lysosomal enzyme responsible for degrading ceramides. Homozygous loss-of-function mutations in *GBA1* lead to Gaucher disease, whereas a mutation in one allele can result in Parkinson’s disease with an odds ratio of 5.4^36^. To better understand which genes in these sgRNA libraries regulate lysosomal lipid degradation we performed a screen for GCase activity by treating cells with PFD-FDGlu, a ceramide linked to a fluorophore and quencher. Upon cleavage of the ceramide by GCase, the fluorophore is de-quenched leading to increased fluorescence. As expected, KD of GBA1 by CRISPRi resulted in reduced GCase activity in our screen (Figure 2J), and increased GBA1 expression by CRISPRa increased GCase activity (Figure 2K). In addition to *GBA1*, *LDLR*, *AXL*, *TREM2, CD81* and *LILRB4* were bidirectional regulators of GCase activity, except that *LILRB4* and *CD81* showed the opposite directionality compared to the other genes. To further characterize the lysosomal phenotype, we conducted additional screens measuring lysotracker signal and sensitivity to lysosomal stressors. The previously mentioned genes were not identified as hits in these control screens, suggesting their genetic manipulation affects lysosomal function rather than abundance (Figures S2D–L).

Next, we validated some of the hits in an additional myeloid cell type. We used primary human peripheral blood mononuclear cell (PBMC)-derived macrophages (hMΦ) so that we could capture donor-to-donor variance in our validation studies. We focused on validating top hits from the GCase screen because this assay was most directly relevant to patients with disease. Because macrophages are difficult to transduce with lentivirus^37^, which is commonly used to deliver dCas9 and sgRNAs, we used electroporation of Cas9 ribonucleoproteins (RNPs) containing either non-targeting control sgRNAs or sgRNAs against candidate target genes to validate the CRISPRi screen hits. For each gene, we chose 2 sgRNAs ∼20–50 bases apart from each other and used Sanger sequencing to confirm KO of the target gene (Supplementary Table 3). As expected, *GBA1* KO led to strongly diminished GCase activity consistently across hMΦ from 3 donors (Figure 2M). This reduction in GCase activity was similar in scale to the activity seen in control cells treated with Conduritol B Epoxide (CBE), a potent GCase inhibitor (Figure 2M). KO in hMΦ of *SPI1*, *TNFRSF1A*, *CLIC1*, and *BIN1* but not *LDLR* also resulted in the expected GCase phenotype based on the screening results, suggesting that most of the genes identified in the THP-1 screens may validate in additional myeloid cell types (Figure 2M, N).

To validate a hit from our CRISPRa screen, we took advantage of a previously developed TREM2-activating antibody^17,38^ and used it to activate TREM2 signaling in hMΦ. While loss-of-function mutations in TREM2 have previously been shown to reduce lysosome function^39^, consistent with our CRISPRi screen (Figure 2J), our CRISPRa screen suggests increasing TREM2 signaling in WT cells may result in increased lysosome function (Figure 2K). To test this, we treated hMΦ with a TREM2 activating antibody, AL002, or an isotype control antibody and then measured GCase activity 3 days later. AL002 led to increased GCase activity in hMΦ from 4 donors (Figure 2O). Interestingly, despite the differences in the distribution of GCase activity from isotype-treated macrophages from different individuals (Figure 2P, gray), AL002 consistently shifted the population of cells toward higher GCase activity (Figure 2P, colors). Taken together, our screens identified a core set of genes that altered lipid uptake and lysosome function, often bidirectionally in THP-1 cells (Figure 2Q).

As a first step toward integrating the screen data, we used hierarchical clustering to cluster the screen data, both across the perturbations and across the functional assays. As expected, in the CRISPRi data, the two lipid uptake screens, APOE and LDL uptake, were closely correlated, as were the two lysosome function screens, lysosome proteolysis and GCase activity (Figure S3A). Interestingly, all of the lysosome and lipid screens clustered more closely with each other than they did with the surface TREM2 screen, which clustered closely with the survival screen, consistent with TREM2’s known role in survival and growth^15–17^. Probably the most striking finding from this clustering analysis though, was that when grouping genes based on their performance across CRISPRi screens, TREM2 clustered closely with its co-receptor TYROBP (Figure S3A). This suggests that performing screens focused on different aspects of cell biology and then integrating the data may contribute to large-throughput unbiased functional annotation of genes, one of the major challenges of the genomics era ^40,41^.

### Perturb-seq in primary human macrophages reveals transcriptomic impacts of CRISPRa/i perturbations

Myeloid cells, including macrophages and microglia, obtain different subtypes (e.g. M1 vs M2, or homeostatic vs disease-associated, respectively) in response to extracellular cues. In AD, these subtypes are primarily defined by transcriptomic signatures. To gain insights into how the genes in our custom sgRNA library may alter the transcriptomic signature we developed a strategy for performing Perturb-seq, a technique that combines pooled CRISPR screening with single-cell RNA sequencing. We used hMΦ because they are a better surrogate for *in vivo* human microglia, and we thought hMΦ would be more capable of obtaining different cell subtypes than the THP-1 cells used in the above screens. Briefly, we infected hMΦ with a pool of lentiviral constructs that encode our custom sgRNA libraries and an RFP reporter. At the same time, we also infected the cells with adeno viruses containing dCas9-VPH or dCas9-KRAB and a GFP reporter. One week after transduction we sorted the cells with FACS to select those cells infected with both viruses and performed scRNA-seq (Figure 3A).

**Figure 3:**
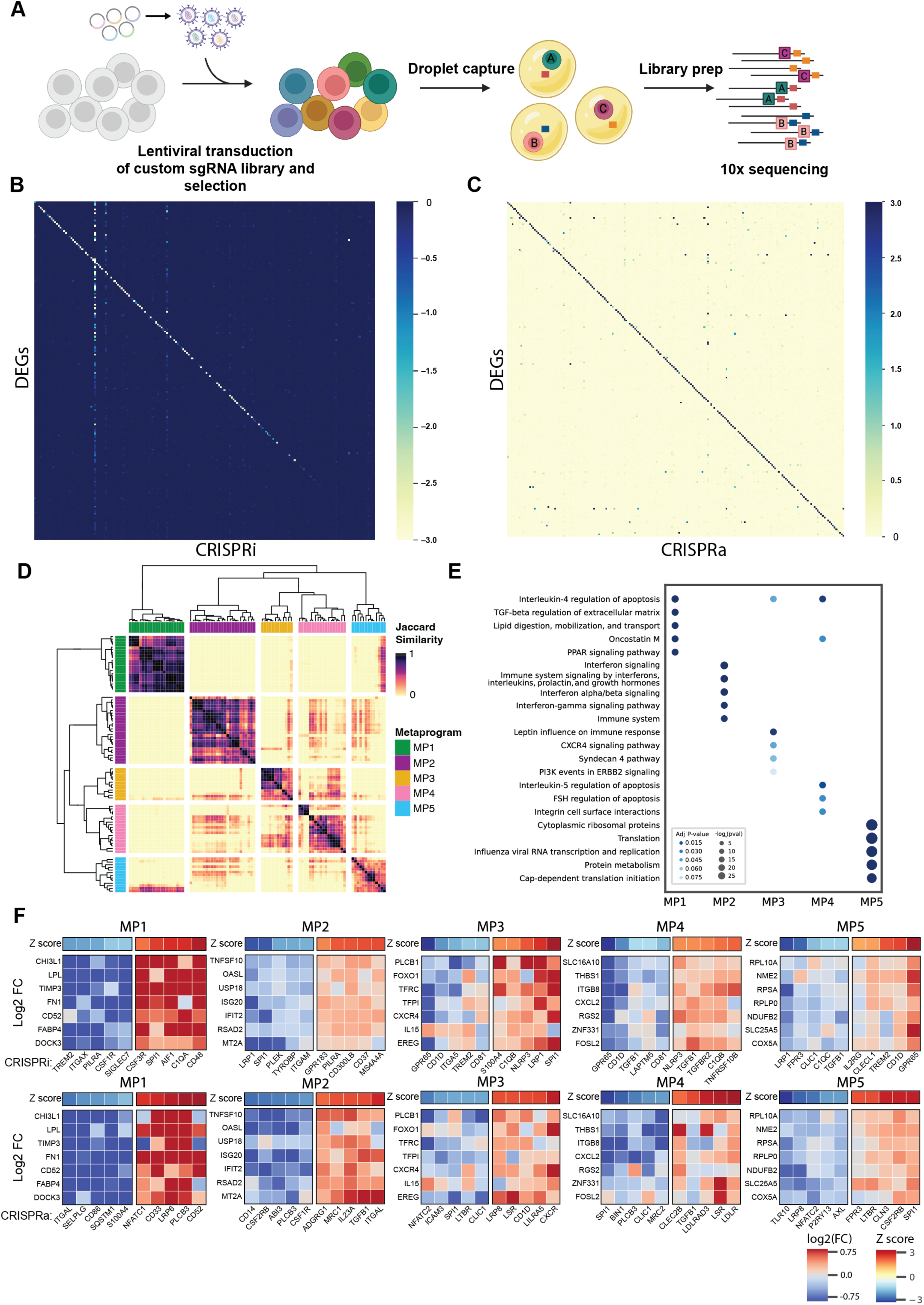
Perturb-seq in primary human macrophages via CRISPRa/i. A) Schematic depicting the experimental workflow for the Perturb-seq experiment. B and C) A matrix of RNA seq results for CRISPRi (B) or CRISPRa (C) filtered to only look at the expression of the genes in the library. Columns represent a particular target gene, whereas rows represent the resulting expression of targeting that gene. Genes are ordered from top to bottom based on baseline expression in non-targeting controls. Color represents the signed log2 fold change (log2FC) values multiplied by the –log10 Benjamini-Hochberg (BH) corrected p-values, relative to non-targeting controls. The visible diagonal in both plots represent on-target efficiency. D) Expression values from CRISPRi and CRISPRa Perturb-Seq experiments were used to generate a pairwise similarity matrix (Jaccard index) of gene programs that comprise each metaprogram, clustered with hierarchical clustering. E) Enrichment analysis of genes within each metaprogram using the mSigDB Hallmark gene set library. The size and color of the dots represent p-values from Enrichr. F) Heatmaps of representative genes in each metaprogram. The values displayed in each heatmap are log2FC compared to NT controls for top 5 activators and repressors of each program. The top bar represents Z-transformed average module scores for each metaprogram.

To assign guides to cells, for each library, we systematically assessed the lowest unique molecular identifier (UMI) threshold that could accurately make assignments, then we confirmed accuracy through differential expression analysis with *glmGamPoi*^42^ using the hMΦ donor as a covariate to correct for the donor effect (Figure S4A–E). 33.7% of cells were assigned to a single guide for CRISPRi and 42.9% for CRISPRa (Figure S4A). For cells containing more than one sgRNA targeting different genes, we assigned the guide with the highest UMI resulting in 28,446 cells for CRISPRi and 23,509 cells for CRISPRa. This filtering criteria resulted in a median coverage of 123 cells/gene at a mean read depth of 8879 reads/cell for CRISPRi, and 106 cells/gene at a read depth of 9657 reads/cell (Figures S3A, S3B).

As expected, CRISPRi guides were more effective at on-target knockdown for genes that had higher expression (Figure 3B). For CRISPRa, there was not a clear connection between guide efficacy and basal expression (Figure 3C). Relative to the non-targeting (NT) controls, we observed a median target knockdown of 57.4% (log2 fold change = −1.228) with the CRISPRi library and overexpression of 289% (log2 fold change 1.950) with the CRISPRa library (Figure S4C). This confirmed both the efficacy of our CRISPR libraries and sgRNA assignment strategy. For all further downstream analyses, we excluded perturbations in which the target gene was not significantly affected by its respective guides (identified through *glmGamPoi* DEG analysis), leaving 86 genes for CRISPRi and 172 genes for CRISPRa.

### Identification of robust gene modules across CRISPRa/i

To reduce dimensionality of the Perturb-seq dataset we identified groups of genes co-regulated by the perturbations. We used non-negative matrix factorization across both CRISPR modalities and both hMΦ donors using *GeneNMF* to identify metaprograms (MPs), representing different cell states^43^. We filtered out MPs if they had a sample coverage of less than 100%, a silhouette score that was less than 0.15, or were comprised of fewer than 5 genes. This approach retained 5 MPs across both CRISPR modalities (Figures 3D). Using the Bioplanet 2019 gene set library, we used *Enrichr* to uncover the functions associated with the MPs^44–46^ (Figure 3E). Hierarchical clustering of the MPs showed that MP2, MP3, and MP4 were closely related, likely due to their roles in various aspects of stress response (Figure 3D, E). Overall, we identified five distinct MPs that are defined by expression of different sets of genes representing different cell states.

To identify which genes in the custom sgRNA libraries contribute to different cell states we calculated MP signature scores for each cell using Seurat’s *AddModuleScore*. We grouped cells that received guides targeting the same gene and calculated an average MP signature score for each MP. Then we performed a Z-score transformation on the average signature scores. To visualize the impact of a perturbation on a MP we plotted the Z-score or the Log_2_ FC of the component genes of a MP (Figure 3F, S5A, B, Supplementary Table 4). Overall, CRISPR perturbations altering a particular MP Z-score altered most of the component genes from that MP in the same direction, confirming this approach (Figure 3F, S5A, B, Supplementary Table 4).

MP1 was enriched in genes involved in extracellular matrix remodeling and lipid metabolism, including *FABP4*, *TIMP3*, *LPL,* and *CHI3L1,* which encodes the AD biomarker YKL-40^47^ (Figure 3E, F, S4F, S5A, B, Supplementary Table 4). CRISPRi-mediated reduction of TREM2 emerged as the strongest regulator of these genes, consistent with TREM2’s known role in lipid homeostasis^13^ (Figure 3F, S4F, S5A, Supplementary Table 4). *CSF1R*, another vital gene for microglia survival, proliferation, and phagocytosis, also reduced MP1 expression when reduced with CRISPRi (Figure 3F, S4F, S5A, Supplementary Table 4). MP2 contained genes tied to interferon signaling such as *IFIT2*, *ISG20*, and *TNFSF10* (Figure 3E, F, S4F, S5A B, Supplementary Table 4). Reduction of PILRA, which upregulated MP1 genes, led to decreased expression of MP2 genes (Figure 3F, S4F, S5A, Supplementary Table 4), suggesting opposing transcriptional regulation between MP1 and MP2. Consistent with this, reduced *SPI1* expression, which upregulated MP1 genes, decreased MP2 gene expression (Figure 3F, S4F, S5A, Supplementary Table 4).

MP3 was enriched in genes linked to immune response, including *CXCR4*, *OLR1* and *IL15* (Figure 3E, F, S4F, S5A, B, Supplementary Table 4). *SPI1* was a bidirectional regulator of MP3 genes; its reduction increased MP3 gene expression, whereas its activation suppressed MP3 gene expression (Figure 3F, S4F, S5A, B, Supplementary Table 4), consistent with *SPI1*’s known role in regulating the inflammatory response of microglia and macrophages^48^. Clustering analysis showed that MP3 and MP4 were the most closely related MPs to each other (Figure 3D). As expected from their relatedness, MP3 and MP4 genes were similarly altered by the same genetic perturbations. For example, reduced GPR65, CD1D, or CD81 lowered expression of both MP3 and MP4 genes, whereas reduced expression of C1QB and NLRP3 increased expression of both MP3 and MP4 genes (Figure 3F, S4F, S5A, Supplementary Table 4). Similarly, overexpression of CLIC1 or LSR decreased or increased, respectively, both MP3 and MP4 genes (Figure 3F, S4F, S5B, Supplementary Table 4). MP5 was composed primarily of ribosomal proteins (RPL10A, RPSA, RPLP0, ext.) and mitochondrial proteins (NDUFB2, SLC25A5, COX5A, ext.) (Figure 3E, F, S4F, S5A, B, Supplementary Table 4). Upregulation of genes encoding these organelles can occur during proliferation or activation^49,50^. Either decreasing *TREM2* expression with CRISPRi or increasing *SPI1* expression with CRISPRa led to enhanced MP5 expression (Figure 3F, S4F, S5A, B, Supplementary Table 4).

Interestingly, in the Perturb-seq experiment, manipulating *SPI1* and *TREM2* expression consistently pushed the cells in opposite directions. For example, reduced expression of *TREM2* increased expression of MP1, MP3, and decreased expression of MP5 genes, whereas reduced expression of *SPI1* decreased MP1 and MP3 genes (Figure 3F, S4F, S5A, B, Supplementary Table 4). Similarly, increasing expression of *SPI1* and reducing expression of *TREM2* both resulted in decreased MP3 genes and increased MP5 genes (Figure 3F, S4F, S5A, Supplementary Table 4).

### Perturbation of *SPI1* has significant downstream effects in human macrophages

Since we used the same libraries for functional assays as we did for Perturb-seq we could identify genetic manipulations that affected both the transcriptome and functional outcomes. To assess the impact across multiple assays, we computed a composite score by calculating the sum of the absolute values of Z-scores in the functional assays and plotted it against the number of differentially expressed genes (DEGs). A few genes were top hits across functional assays but didn’t have many DEGs, or vice versa, but by-in-large, most of the top hits had both a transcriptional and functional phenotype (Figure 4A). For example, *SPI1*, which encodes PU.1, a critical myeloid cell transcription factor that is associated with AD through GWAS ^51,52^, was one of the top hits across the lipid and lysosome screens and the Perturb-seq experiments (Figure 4A). CRISPRi of *SPI1* resulted in a 67.6% reduction in *SPI1* expression (log2 fold change of −1.637) causing 444 upregulated and 392 downregulated DEGs (BH adj p < 0.05 and log2FC > 0.1 or < −0.1), whereas CRISPRa resulted in 34 upregulated and 14 downregulated DEGs (BH adj p < 0.05 and log2FC > 0.1 or < −0.1) (Figures 4B, C). GSEA using the MSigDB hallmark gene set revealed that the DEGs from CRISPRi are involved in IL2/STAT5 signaling, PI3K/Akt/mTOR signaling, and interferon gamma response (Figure 4D). Although fewer DEGs were observed with CRISPRa, *SPI1* activation still significantly affected several important pathways such as TNFα signaling and hedgehog signaling, which is a vital response for macrophage polarization and metabolism (Figure 4E). Remarkably, nearly half (21/48) of the DEGs altered by CRISPRa of *SPI1* were regulated in the opposite direction by CRISPRi of *SPI1*, suggesting those genes may be more directly regulated by *SPI1* expression levels when compared to the 835 DEGs regulated by reduced *SPI1* levels with CRISPRi (Figure S6A, B). Overall, these results align well with previous findings on the function *SPI1*, further validating the reliability of our Perturb-seq results^52–54^.

**Figure 4:**
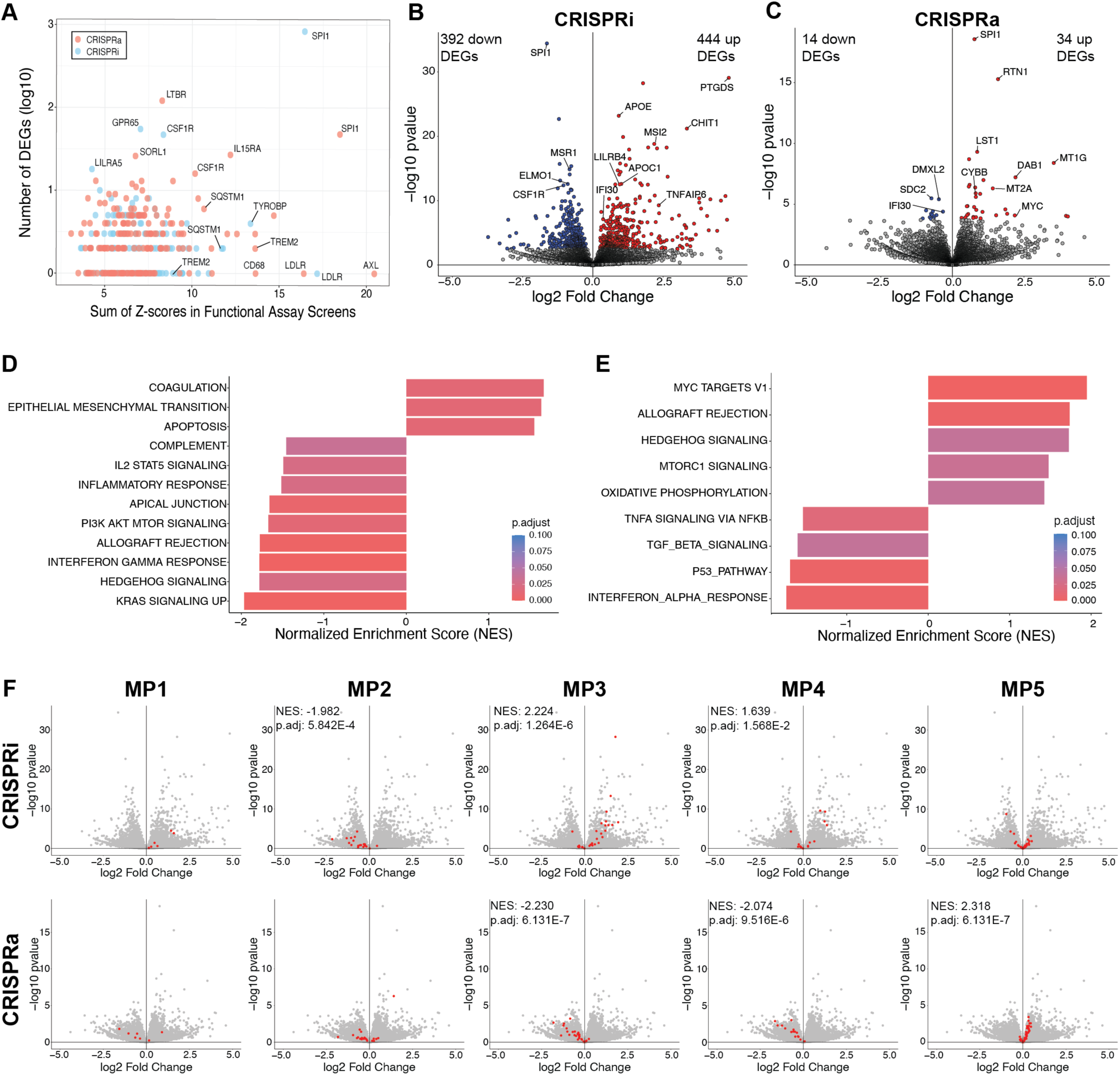
Downstream transcriptional effects of SPI1/PU.1 knockdown and overexpression. A) Dotplot depicting the impact of CRISPR perturbations across assays for CRISPRa (red) and CRISPRi (blue). The x-axis represents a composite score, calculated by the sum of the absolute values of Z-scores from all functional assays. The y-axis indicates the number of differentially expressed genes (DEGs) with a p-value < 0.05 for each perturbation compared to NT controls. B and C) Volcano plots showing downstream effects of CRISPRi (B) and CRISPRa (C) perturbation of SPI1 compared to NT controls. DEGs (p-value < 0.05 and |log2FC| > 0.1) are colored. D and E) Gene set enrichment analysis of upregulated and downregulated DEGs for SPI1 inhibition (D) and SPI1 activation (E) using the mSigDB Hallmark gene set library. The x-axis shows normalized enrichment scores, and the bars are colored by adjusted p-values. F) Volcano plots for CRISPRi (top row) and CRISPRa (bottom row) of SPI1 perturbation overlaid with genes in each MP (highlighted in red). Significantly enriched MPs denoted with normalized enrichment scores (NES) and adjusted p-values.

We then looked at the distribution of the component genes from the MPs within either the CRISPRi or CRISPRa *SPI1* volcano plots (Figure 4F). Overall, as expected, component genes of a MP moved in the same direction when that MP was altered by *SPI1* expression. For example, CRISPRi of *SPI1* was the strongest regulator of MP3 genes (Figure 3F, Figure S5A, Supplementary Table 4), and most MP3 genes were increased by CRISPRi of *SPI1* (Figure 4F). Interestingly, *SPI1* perturbation showed significant bidirectional effects on MP3 and MP4 (Figure 4F), consistent with these two programs being most closely related to each other (Figure 3D). Although not significant, a similar trend occurred for MP1, genes in MP1 were mostly upregulated with *SPI1* inhibition and downregulated with activation (Figure 4F).

### Highly correlated genes across all assays are more likely to be functionally linked

In total, we generated 15 CRISPRa and 15 CRISPRi screen datasets, consisting of 3 lipid assays, 5 lysosomal assays, surface TREM2, cell growth, and the 5 MPs identified through Perturb-seq (Figure 5A). Initially, we assessed the function of the MPs using *Enrichr* (Figure 3E). As an independent strategy to assess the relationships between the functional assays and MPs we used Pearson correlation to compute pairwise correlations using the CRISPRi perturbation response matrix (Figure 5B). By calculating correlation coefficients across assays, we identified patterns of similarity, allowing us to determine which assays yielded similar responses to perturbations. Like before when this analysis was done without inclusion of the MPs (S3A), in the CRISPRi screens, the two lysosomal stressor screens clustered together, as did the two lipid uptake screens, confirming this approach. Interestingly, MP5, which consisted mostly of ribosomal, mitochondrial, and other housekeeping genes, clustered closely with the Bodipy screen, consistent with a known crosstalk between the proteostasis network and lipid homeostasis^55^. MP1, which *Enrichr* showed was enriched in lipid digestion genes (Figure 3E), clustered with the Lysotracker screen (Figure 5B), consistent with the lysosome being the site of lipid digestion after endocytosis^28^.

**Figure 5:**
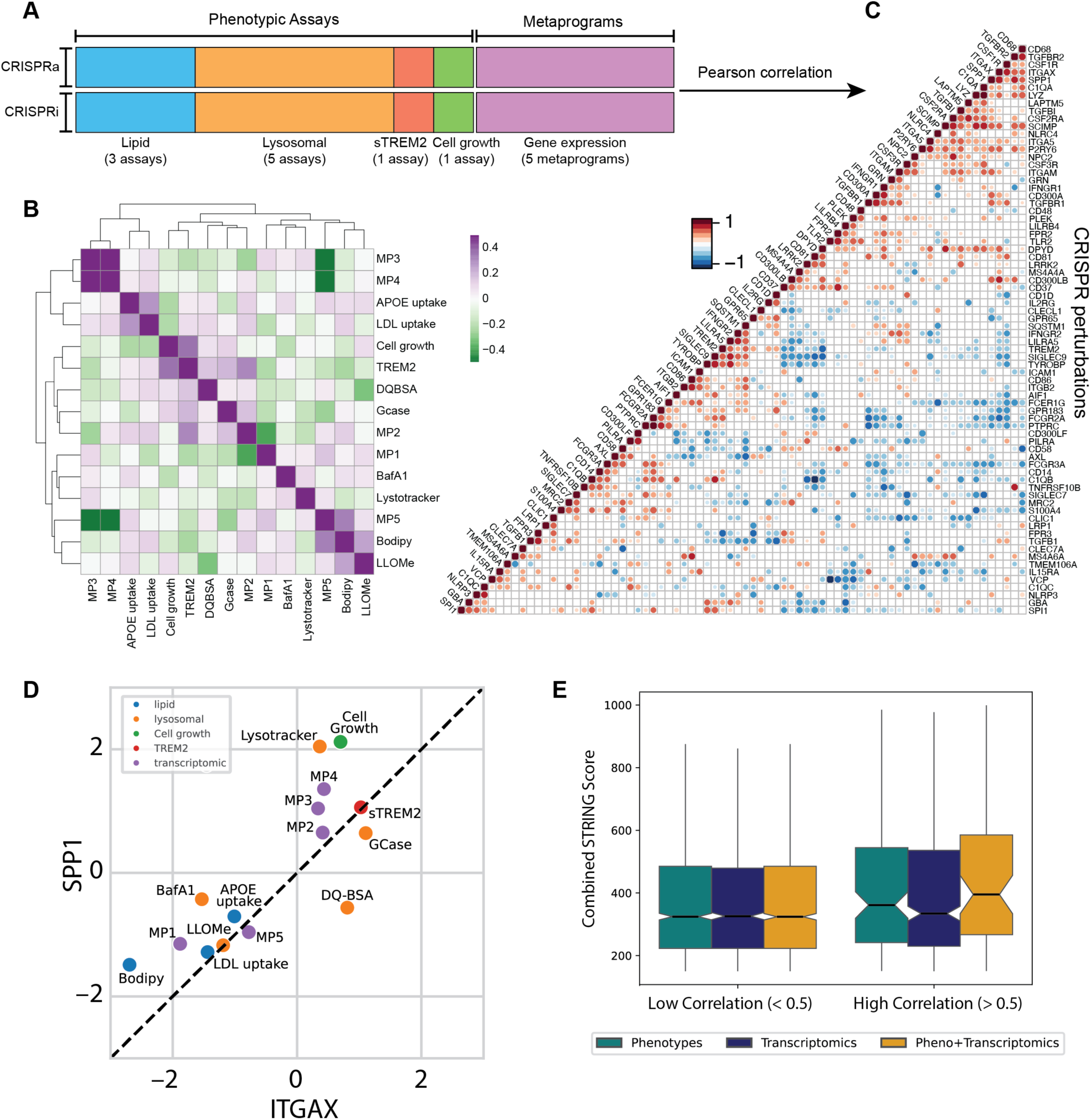
Integrated analysis using phenotypic assay and transcriptomic data uncovers gene pairs with similar functional roles. A) Schematic summarizing the screen data collected in this study. Assays are colored by functional class; transcriptomic data is summarized into metaprograms identified in Figure 3. B) Correlation matrix showing the relatedness of CRISPRi screen results using Pearson correlations. Positive correlations are shown in purple and negative correlations in green. C) Gene-gene correlation matrix computed using Pearson correlation across all CRISPRi genes. Ward.D2 linkage method was used to perform hierarchical clustering. Dot size within the matrix represents the p-values calculated using cor.mtest with a 95% confidence interval and dot color corresponds to the Pearson correlation coefficient. Non-significant (p < 0.05) pairs are shown as blanks. D) Dot plot showing the Z-score of ITGAX (x-axis) and SPP1 (y-axis) in the individual screens. Dots are colored according to the functional classes of the assays defined in (A). E) Comparison of STRING interaction scores for gene-gene pairs in the CRISPRi library grouped based on correlation strength, classified by highly correlated (r >= 0.5) or weakly correlated (r < 0.5). STRING interaction scores were calculated for gene-gene pairs defined using the combined matrix from (A) (orange), only the functional assay data (green), or the metaprogram transcriptomics alone (dark blue).

To evaluate which individual gene perturbations perform similarly across the assays screened here, we focused on genes that were (1) significantly perturbed in the Perturb-seq and (2) showed at least one strong phenotype (|Z-score| > 1.645) across these assays (Figure 5A). For CRISPRi, this filtering criteria left 74 genes. Using transformed Z-scores for each metaprogram and the normalized Z-scores from the functional assays, we created a combined matrix (Figure 5C). *SPP1*, and *ITGAX*, which encodes CD11c were highly correlated genes across these assays with a Pearson correlation of 0.791 (Figure 5D). CRISPRi of either gene resulted in reduced ApoE4 and LDL uptake, reduced expression of MP1 and MP5 genes, decreased lipid levels, increased sensitivity to the lysosome stressors Bafilomycin A1 and LLOME, increased cell growth, increased lysosome content and GCase activity, increased surface TREM2, and increased expression of MP2, MP3, and MP4 genes.

This combined data matrix is a unique resource. It enables gene product function prediction of uncharacterized genes based on the genes clustering with better characterized genes across the combined data matrix. To evaluate the validity of this approach we used the STRING database, which uses genomic context, gene co-expression, interactome experiments, manually curated resources, and text mining to quantify the likelihood that two proteins interact, either physically or together in the same pathway^56,57^. Using the 71 genes remaining in the CRISPRi library after filtering we obtained the STRING score for every possible gene pair. Then we calculated a combined STRING score for either the gene-pairs with a low correlation in our assays (Figure 5A, C, E) or a high correlation in our assays (Figure 5A, C, E). CRISPRi gene pairs with higher correlation in our assays (R>=0.5) showed a higher STRING protein-protein interaction confidence score (Figure 5E). If we compute these pairwise correlations using only the functional assay data or only the transcriptional metaprogram data, we see a lower average STRING interaction confidence score, demonstrating the power of combining transcriptomic datasets with cellular assay-based screening.

## Discussion

In this study, we generated custom targeted CRISPRa/i sgRNA libraries and used them to identify mediators of lipid homeostasis, lysosomal function, and surface TREM2. Using targeted libraries for these studies enabled screening at higher than typical coverage (usually ∼8,000 cells screened/sgRNA in the library), and it enabled us to perform 10 different functional screens for each of CRISPRa and CRISPRi in the myeloid model, THP-1 cells. Whereas most CRISPR screening studies focus on a single CRISPR modality of one functional assay, this unique dataset enabled a more comprehensive mechanistic understanding of the targeted genes.

Although several microglia genes are well established genetic risk factors for AD, there are no microglia-targeting therapeutics approved yet. Direct TREM2 activation with two different TREM2-agonist antibodies did not show beneficial effects in Phase 2 clinical trials for AD^58^ or the neurodegenerative disease adult-onset leukoencephalopathy with axonal spheroids and pigmented glia (ALSP) caused by CSF1R mutations^59^, while a small molecule agonist is still being investigated in clinical trials^19^. An alternative or complementary strategy to activating TREM2 directly is to increase cell surface TREM2 levels, priming microglia for activation when they encounter a TREM2 ligand. This approach may result in only those microglia that are both primed by the therapeutic and locally triggered by pathology to become activated, in contrast to globally activating TREM2 in the brain. Such an approach could be particularly beneficial when paired with anti-Aβ therapeutics. In our study, we identify TGFBR2 as a key regulator of surface TREM2 levels in THP-1 cells, establishing a functional link between two AD genetic risk loci^30^ and highlighting TGFBR2 as a potential therapeutic target for combination therapy with anti-Aβ antibodies in AD.

One limitation of using THP-1 cells as a screening model is their low expression of some critical microglial genes including MS4A6A. We previously demonstrated that in primary human macrophages and iPSC-derived microglia, degradation or knockdown of MS4A4A leads to stabilization of TREM2 in the cell membrane in a MS4A6A- and DAP12-dependent manner^60^. Consistent with these findings, we show here that DAP12 (TYROBP) expression bidirectionally regulates membrane TREM2 levels in THP-1 cells. In contrast, MS4A4A overexpression failed to decrease membrane TREM2 levels, likely due to the low expression levels of MS4A6A, a key component of this signaling pathway. Instead, in THP-1 cells, MS4A4A overexpression was a main driver of membrane TREM2 levels.

These results underscore the importance of cellular context in interpreting screen outcomes and highlight a broader advantage of performing both loss-of-function (LOF) and gain-of-function (GOF) screens. By leveraging both LOF and GOF screens, we highlighted genes that bidirectionally regulate a phenotype, which are generally more likely to be key regulators of a pathway. However, overreliance on this approach can be misleading as there are many biological reasons why genetic manipulation in only one direction may cause a phenotype. For example, in some cases, reducing expression of a gene encoding one peptide that is part of a heteromultimeric protein complex may disrupt the complex, whereas overexpression of only one component would not increase expression of the full complex. Another benefit to performing both LOF and GOF screens is that for many human genetics-implicated genes, the directionality of the mutation is not clear. Alternatively, sometimes human genetic data does clearly support one direction being detrimental, raising the possibility that the opposite direction may be beneficial. Performing both GOF and LOF screens can help clarify directionality of disease mutations and evaluate whether the opposite direction of a disease driver could be beneficial in a wild-type background.

In addition to performing paired bidirectional screening, this dataset is unique in the number of screens performed. We propose that integrating data across multiple screens contributes to deeper mechanistic understanding of gene function. For example, it’s interesting that several genes including *TREM2*, *AXL*, *SCARB1*, and *ABCA1* emerged as top regulators in both lipid uptake and lysosome function screens, consistent with the inter-relatedness of these pathways. Notably, these genes were not hits in the Lysotracker or steady-state lipid screens, suggesting a more direct regulatory role in these pathways. In contrast, LDLR reduction with CRISPRi reduced lysosome abundance in the cells (Figure S2G), leading to reduced lysosome function in both the lysosomal proteolysis screen (Figure 2G) and reduced GCase activity (Figure 2J).

However, this phenotype was not replicated in the GCase assay performed on primary macrophages, suggesting there may be cell-type specific differences in LDLR regulation of lysosomal abundance in primary macrophages and THP-1 cells (Figure 2M). These detailed insights are made possible by combining screen data from many different cellular assays.

Integrating the data from many screens is a powerful approach, but it’s biased by the types of screens performed. In contrast, transcriptomics enables unbiased insights into the molecular function of the targeted genes. We performed Perturb-Seq studies to determine the transcriptomic signature of the genes in the targeted libraries. However, limited sequencing depth is a common problem in Perturb-seq studies, owing to the high cost of reagents and sequencing^61^. To overcome this limitation, we used unbiased approaches to define 5 MPs, which represent different cell states. This approach provided increased power to detect sgRNA-driven transcriptional changes. Integrating the transcriptional data with the functional assay data enabled two unique analyses not commonly done in CRISPR screen studies. First, at the screen level, we could gain a deeper and unbiased understanding of the MPs themselves. For example, clustering of the screen data allows for determining which MP expression is regulated by the same genes that regulate the functional assays included in this study. Second, at the gene level, we can identify genes whose manipulation results in a similar phenotype across the assays screened here. This approach is a unique and scalable strategy for unbiased characterization of poorly characterized genes, a problem that has remained a challenge throughout the genomic era. While the current study was limited to ∼200 genes in a targeted library and 10 CRISPRi functional assays and 5 MPs, future studies could be scaled to thousands of genes.

In summary, this study not only identifies key regulators of lipid homeostasis, lysosomal function, and microglial activation, but also provides a multidimensional resource for the field. By integrating CRISPRa/i functional screens with transcriptomic profiling, we create a scalable framework for systematic gene function discovery.

The resulting dataset—linking genetic perturbations to cellular phenotypes and transcriptional programs— serves as a valuable resource for exploring microglial gene function in neurodegenerative disease and beyond.

## Methods

### Cell culture

THP-1 cells (ATCC, #TIB-202) were maintained in myeloid media (RPMI medium (Corning, #10-040-CV) supplemented with 10% FBS (Fisher Scientific, #SH3007103HI) and 1% penicillin/streptomycin (Thermo Fisher, #15140122), 1% MEM NEAA (Thermo Fisher, #11140-050), Glutamax (Thermo Fisher, #35050061), and sodium pyruvate (Thermo Fisher, #11360070)). Cells were diluted in fresh medium every 3–4 days at a density of 0.2 million cells/mL and replated on non-TC-coated tissue culture plates. Stable tetracycline-inducible CRISPRi or CRISPRa THP-1 lines were generated with lentivirus from plasmids in which Cas9 from Cellecta #SVRTC9E2B-PS was replaced with either dCas9-KRAB or dCas9-VPH from Cellecta #SVKRABC9SFB-PS or Cellecta #SVVPHC9SFB-PS, respectively. 2 days after transduction, cells were grown in the presence of 10 µg/mL Blasticidin for 1 week, expanded and banked. Cells were then transduced with sgRNA libraries at an MOI of 0.3, selected with 1 µg/mL puromycin for 1 week, expanded and banked.

Primary human monocytes were isolated by adding 50 µL 0.5 M EDTA (Thermo Fisher, #AM9260G) to 25 mL whole blood (Vitalant), incubating with 1.25 mL RosetteSep Human Monocyte Enrichment Cocktail (Stem Cell Tech., #15068) for 30 min at room temp, and diluting with 25 mL PBS containing 2% FBS and 1 mM EDTA. Sepmate tubes (StemCell Tech., #85460) were prepared by adding 14 mL Ficoll-Paque Plus (Cytiva, #171404003) to the tubes, centrifuged at 300 x g for 2 min to remove bubbles, then loaded with the previously prepared blood and centrifuged at 1200 x *g* for 20 min at room temp using a slow break to stop. Monocytes were collected, washed in PBS containing 2 % FBS and 1 mM EDTA, and residual red blood cells were lysed by adding 2.5 mL ACK lysis buffer (Thermo Fisher, #A1049201) to the pellet. The cells were washed again in PBS containing 2 % FBS and 1 mM EDTA, and resuspended and plated in macrophage media (myeloid media supplemented with 100 ng/mL of M-CSF (Biolegend #574808)). After 6 days, macrophages were harvested by removing media, incubating in PBS for 5 min, then mechanically detached using a cell scraper and subjected to the described treatment.

### Lentivirus generation

293-T cells (Thermo Fisher, #R70007) were maintained in DMEM (Thermo Fisher, #11965092) supplemented with 10% FBS and 1% penicillin/streptomycin, and replated 2–3 times/week by dissociating the cells using TrpLE (Thermo Fisher, #12604013) and replating at a ratio of 1:5 – 1:20. The day before transfection, cells were replated at a ratio of 1:2 so they were ∼90% confluent on the day of transfection. Cells were transfected with Lipofectamine 3000 (Thermo Fisher, #L3000015) by combining 0.674 ug VSV-g, 1.011 ug psPax, 1.347 ug dCas9 transfer plasmid, and 7.415 µL P3000 Reagent in 100 µL Opti-MEM (Thermo Fisher, #31985-062) with 7.415 µL of Lipofectamine 3000 in 100 µL Opti-MEM, vortexing briefly, incubating for 10 min at room temp, and adding the solution to 1 well of a 6-well plate of 293T cells.

### sgRNA library design

Using the Human GRCH38 (Ref Seq v.109.20210514) reference genome, CRISPick (https://portals.broadinstitute.org/gppx/crispick/public) was used to select 15 sgRNAs/gene. The gene name and sgRNA sequences were extracted from the CRISPpick output and 5’-ACCG was added to the 5’ end of each sgRNA and 5’-GTTT was added to the 3’ end of the sgRNAs. The sgRNA list was filtered to remove sgRNAs containing a BbsI restriction enzyme site, which would be cleaved during cloning, stretches of 4 or more Ts, which is an RNA pol III termination signal, or those sgRNAs that start with 5 or more Gs, which reduces sgRNA expression. The top 5 remaining sgRNAs for each gene were selected. 100 non-targeting sgRNAs were selected from the Calabrese library^62^ and the library was cloned into a custom expression construct containing a sgRNA scaffold driven by a U6 promotor and containing capture sequence 2 at the 3’ end of the scaffold for use with single-cell RNA sequencing. This construct also contained a CMV promotor-driven RFP expression cassette followed by a P2A-self cleaving peptide sequence then a puromycin resistance gene.

### THP-1 screens

THP-1 CRISPRa or CRISPRi lines containing sgRNA libraries were grown in the presence of 1 µg/mL doxycycline for 7 days to induce dCas9 expression. In some cases, cells were differentiated by growing 25 million cells in 25 mL full RPMI medium containing 1 µg/mL doxycycline and 25 nM PMA for 3 days on a TC-coated T150 flask, then removing PMA for 1 d by changing media to full RPMI medium containing 1 µg/mL doxycycline. Fluorescent-based screens were performed by staining cells as described below and using FACS to sort the top and bottom 10% of cells based on fluorescent signal and isolating gDNA for sgRNA quantification. Survival-based screens were performed by growing the cells in a challenge or control condition and quantifying sgRNA abundance in gDNA from the resulting populations.

GCase screens were performed by treating differentiated THP-1 cells with 150 µM PFB-FDGlu (Thermo Fisher Scientific, #P11947) for 1 h at 37 C, washing 2x with 4 C FACS buffer (Fisher Scientific, #BDB554656), mechanically detaching the cells and sorting on a FACS. Lysotracker screens were performed by treating differentiated THP-1 cells with 100 nM Lysotracker (Thermo Fisher, #L7526) for 5 min at 37 C, washing 2x with 4 C FACS buffer, mechanically detaching the cells and sorting on a FACS. Lysosomal proteolysis screens were performed by treating 10 million THP-1 cells in 10 mL myeloid medium for 1 h with 25 µg/mL DQ Green BSA (Thermo Fisher, #D12050), washing 2x in 4 C FACS buffer, and sorting on a FACS. LDL uptake screens were performed by treating 10 million THP-1 cells in 10 mL myeloid media with 1 µg/mL pHrodo-Green LDL (Thermo Fisher, #L34355) for 3 h at 37 C. ApoE4 uptake screens were performed by incubating 10 million THP-1 cells in 10 mL myeloid media with 10 µg/mL ph-rhodo labeled (Thermo Fisher, #P35369) ApoE4 (Tonbo Biosciences, #21-9195-U500) for 24 h at 37 C, washing 2x in 4 C FACS buffer, and sorting on a FACS. Surface TREM2 screens were performed by staining 10 million cells in 10 mL FACS buffer containing 20% Fc Binding Inhibitor (Thermo Fisher, #14-9161-73) and 50 µL DL650 conjugated (Thermo Fisher, #84536)-T22 anti-TREM2 antibody (Alector) for 30 m on ice, washing 2x with FACS buffer, and sorting on a FACS. Bodipy screens were performed by treating differentiated THP-1 cells that had been serum starved for 24 h with 2 µM Bodipy (Thermo Fisher, #D3922) for 30 min, washing them in 2x FACS buffer, and sorting them on a FACS. Baseline cell growth screens were performed by growing 10 million THP-1 cells for 7 d in the presence 1 µg/mL doxycycline and comparing the sgRNA read counts to either the abundance of sgRNA in the plasmid library used for virus preparation or to sgRNA abundance in replicate samples at time 0 (without 7-day growth in doxycycline). Lysosomal stressor screens were performed by growing cells in the presence of 10 nM Bafilomycin A1 (Thermo Fisher, #J61835.MCR) for 3 d or 100 µM LLOMe (Thermo Fisher, #125130250) for 3 d, and comparing the resulting sgRNA read counts after sequencing to those from samples grown in the absence of a challenge.

### Sequencing of sgRNAs in functional screens

sgRNA sequences were amplified with 21 cycles (98 C for 10 s, 59 C for 30 s, 72 C for 1 m) of PCR using a reaction mixture containing 1 µg gDNA, 500 nM each of GGW545 and GGW546, and 25 µL of NEBNext Ultra II Q5 Master Mix (NEB, # M0544L) in a volume of 50 µL. Barcodes and Illumina adaptors were added in a second PCR reaction mix containing 7.5 µL of the product above, 500 nM each of barcoded forward (GGW550 N501–510) and reverse primer (GGW551 D701–710), 2.25 µL of a 1:300 dilution of Syber Green (Sigma, #S9430), 37.5 µL NEBNext Ultra II Q5 Master Mix in a volume of 75 µL. A test RT-qPCR reaction (98 C for 10 s, 62 C for 30 s, 72 C for 15 s, 30 cycles) was performed with 25 uL of the reaction mixture and the remaining 50 µL was amplified for the number of cycles needed to reach halfway between maximum and minimum signal in the test RT-qPCR reaction, usually ∼5 cycles. PCR cleanup was performed with SPRIselect (Beckman Coulter, #B23318) following the manufactures protocol for Left Side Size Selection using a 1x sample/bead ratio and eluting in 20 µL water (Corning, #46000CI). Sequencing libraries were quantified using NEBNext Library Quant Kit for Illumina (NEB, #E7630S) in reactions containing 4 µL of standard or sample (1:22,500 dilution) and 16 µL NEBNext Library Quant Master Mix following the manufacturers protocol. Up to 24 samples were combined in equal proportion based on the above quantification and sequenced on a MiSeq using MiSeq Reagent Kit v3 (Illumina, #MS-102-3001), usually resulting in ∼1 million reads/sample. Combined samples were diluted to 4 nM total concentration, 5 µL of the 4 nM library was combined with 5 µL of 0.2 N NaOH and incubated at RT for 5 m, then 990 µL of HT1 (from the Illumina kit) was added. 360 µL of the resulting 20 pM library was combined with 240 µL of HT1 resulting in a 12 pM solution, which was loaded into position 17 of the reagent cartridge. 600 µL of a custom primer (GGW549) for read 1 was prepared at 0.5 µM using HT1 buffer and loaded into position 18 of the reagent cartridge. Sequencing was then initiated on the MiSeq. Reads were trimmed to include only the 20 base sgRNA sequence using Trimmomatic v39, mapped to the sgRNA library using Bowtie v1.3.0, and enrichment analysis was performed with MAGeCK^31^.

### Surface and Soluble TREM2 measurements

THP-1 cells were re-plated at 20,000 cells/well in non-TC 96-well U-bottom plates and treated with vehicle or 20 ng/mL TGF-β1 (R&D Systems, #7754-BH) for 2 days. The cells were pelleted, the media was collected and used to measure soluble TREM2, and the cells were stained in FACS buffer containing 20% FC blocker (Thermo Fisher, #14-9161-73) and 1:50 dilution of DL650-conjugated T22 antibody against TREM2 (Alector) or a DL650-conjugated isotype control (Alector). A custom MSD was used to detect soluble TREM2 in the media. Multi-Array 96 well plates (Mesoscale Discovery Cat #L15XA-3) were coated overnight at 4C with 50 uL of 1 mg/mL anti-TREM2 Clone 8F11 capture antibody (Alector). The next day, plates were washed three times with 3% BSA (P Biomedicals #820451) in PBS for two hours. 50 µL of sample, diluted 1:2–1:10, depending on the experiment, or human TREM2-Fc standard (R&D Systems, #1828-T2-050) were added to the plate and incubated at room temp for 1 h with shaking. Plates were washed 3x in PBS +0.5% Tween, then 50 mL of 100 ng/mL goat anti-human biotinylated TREM2 detection antibody (R&D Systems, #BAF1828) was added. After shaking at room temperature for one hour, plates were washed and 50 mL SulfoTag Strepavidin (MSD #R32AD-1), diluted 1:2,500, was added to the plates. After shaking at room temperature for 30 min, plates were washed and 150 mL of 2X READ Buffer T (MSD, #R92TC-1) was added. Plates were read on a Sector 600MM machine (MesoScale Discovery). Sample concentrations were interpolated from the standard using a 5PL logistic regression model.

### Electroporation of primary macrophages with RNPs

• sgRNAs with a predicted high on-target activity were selected using the Synthego design tool such that the predicted cutsite of each was < 50 bp apart and the deleted fragment from the cutsite predictions was not a multiple of 3 (to minimize in-frame deletions). The sgRNAs were resuspended at 50 µM TE, and 1.65 µL of each sgRNA was combined with 1.05 µL of HF Cas9 (IDT, #1081061) for 10 min at room temp. 15 µL of P3 solution (Lonza, #V4XP3032) was added to the Cas9-sgRNA complex and this solution was used to resuspend 5 million hMΦ that had been pelleted at 300 x *g* for 5 min. The cells were transferred to one well of a 16-channel cuvette (Lonza, # V4XP3032), electroporated using the CM137 program (Lonza), then resuspended in macrophage media, and grown for 1-week on non-TC-coated 6-well dishes, by changing media every 2–3 days. The cells were transferred to U-bottom 96-well plates at a density 50,000 cells/well the day before measuring GCase activity, which was done by adding 150 µM PFB-FDGlu for 1 h at 37 C, washing 2x with 4 C FACS buffer, and quantifying fluorescence using a flow cytometer.

### Perturb seq of primary macrophages

Two donors of hMΦ were each transduced with adenovirus encoding either CRISPRi-IRES-GFP (Vector Biolabs, custom prep) or CRISPRa-IRES-GFP (Vector Biolabs, custom prep), and lentivirus encoding either the CRISPRi or CRISPRa sgRNA libraries, respectively. The cells were grown in macrophage media for 1 week then FACS sorted based on a live cell dye (Thermo Fisher, #C34858), RFP+ (indicating the presence of the guide casette), and GFP+ (marking dCas9 expression). Sorted cells were resuspended in 1X PBS with 0.04% BSA. A total of 20,000 cells per sample were loaded and library preparation was performed according to the manufacturer’s protocol (10x Genomics, CG000206 Rev D) using the 10x Genomics Chromium Single Cell 3’ Gene Expression v3.1 kit with Feature Barcode technology (10x Genomics PN-1000079), generating both gene expression and CRISPR guide capture libraries. cDNA fragment analysis was performed using a Tapestation. Libraries were sequenced using an Illumina NovaSeq X using the following read configuration: Read 1 (10x cell barcode and UMI): 28bp, Read 2 (cDNA): 91bp, and i7 index: 8bp.

### Computational and Statistical Analysis

#### Perturb-seq alignment, cell calling, and guide assignment

Cell Ranger 7.1.0 software (10x Genomics) was used for alignment of scRNA-seq reads to the transcriptome as well as the alignment of sgRNA reads to the library. The *‘cellranger count’* command with default parameters was used to map expression reads to the reference transcriptome (10x Genomics GRCh38 version 2020-A).

Reads from the sgRNA libraries were also mapped using *‘cellranger count’*. A custom script was used for guide assignments. UMI thresholds ranging from 3 and 40 were systematically looped through and guides were assigned to cells based on the UMI threshold. If a cell had multiple guides assigned, the guide with the highest UMI count was retained. For each threshold, differential gene expression analysis was conducted for all perturbations using the python package *glmGamPoi*^42^, with the hMΦ donor as a covariate.

Guide assignments were considered correct if the gene targeted by the assigned guide was differentially expressed (Benjamini-Hochberg corrected p-value < 0.05) and the sign of the log2 fold-change was in the expected direction (ie., negative with CRISPRi and positive with CRISPRa). The final UMI threshold selected for downstream analysis was the threshold that resulted in the highest number of correct assignments, which was 5 UMI for both CRISPRi and CRISPRa. Cell-guide assignments were loaded into a Seurat object (Seurat v5). Cells with poor quality were removed based on mitochondrial gene content and RNA feature counts. For the CRISPRi library, cells with mitochondrial gene percentages above 18%, fewer than 1,000 or more than 7,000 detected features, or total RNA counts exceeding 35,000 were excluded. For the CRISPRa library, cells with mitochondrial gene percentages above 20%, fewer than 1,000 or more than 7,000 detected features, or total RNA counts exceeding 25,000 were filtered out. The resulting matrix was normalized and scaled using the standard steps in Seurat for downstream analysis.

### Identification of metaprograms

Non-negative matrix factorization was performed on the Perturb-seq data using the R package *GeneNMF*^43^. Seurat objects for both CRISPRa/i libraries were filtered to include only genes that were significantly perturbed by their respective guides (Benjamini-Hochberg adjusted p-value < 0.05) and then split by hMΦ donor. Gene programs were identified using the *multiNMF* function across both CRISPR modalities and both donors with parameters set to *k=4:9, min.exp = 0.05*.

Metaprograms (MPs) were identified using the *getMetaPrograms* function with parameters: *metric = “cosine”, weight.explained = 0.8, specificity.weight=7, nMP=8, min.confidence = 0.7.* The resulting MPs were filtered based on the following criteria: (i) sample coverage < 100%, (ii) a silhouette score that was negative or < 0.15, or (iii) were comprised of fewer than five genes. MP identities were determined through enrichment analysis using *Enrichr*^44–46^ with the *Bioplanet 2019* gene set library.

Module scores for each cell were computed using Seurat’s *AddModuleScore* function for each MP. The mean module score was then calculated for each perturbation and standardized to Z-scores using the mean and standard deviation of all module scores for each MP.

### Pathway enrichment analysis of SPI1

Differentially expressed genes (DEGs) associated with SPI1 CRISPRa/i were identified using *glmGamPoi*^42^ with the hMΦ donor as a covariate. Genes were considered to be differentially expressed if the absolute value of the log2 fold change was > 0.1 and the Benjamini-Hochberg adjusted p-value was < 0.05.

To assess pathway-level changes, gene set enrichment analysis (GSEA) was performed using the *GSEA* function from the *clusterProfiler*^63–66^ package. Genes were ranked based on their Z-score and enrichment was assessed using the *MSigDB Hallmark 2020* gene set library. Enrichment analysis of MPs was conducted by constructing a custom gene set library, where each MP was defined by its associated genes. Enrichment scores and significance were then computed using *clusterProfiler’s* GSEA function as mentioned above. Statistical significance was determined using an adjusted p-value threshold of 0.05.

### Correlations of strong perturbations

A combined phenotypic and transcriptomic matrix was created using the transformed Z-scores for each metaprogram and the normalized Z-scores obtained from *MAGeCK*^31^. Only genes that were considered to be “strong perturbations”, i.e., both significantly perturbed in the Perturb-seq (Benjamini-Hochberg adjusted pvalue < 0.05) and exhibiting at least one strong phenotype (|Z-score| > 1.645) across all assays were retained, leaving 74 genes out of the 86 perturbed genes for CRISPRi.

Using the combined matrix, pairwise Pearson correlations were computed for genes based on their perturbation profiles. Correlation matrices were visualized using the *corrplot* package in R^67^ and ordering genes using hierarchical clustering with the Ward.D2 method. To assess statistical significance, a permutation test was performed using the *cor.mtest* function in R with a 95% confidence level. Non-significant correlations (p > 0.05) were removed from the plot.

### STRING interaction scores for gene-gene pairs

Pairwise Pearson correlations were computed to assess relationships between perturbations using (i) the phenotypic matrix only, (ii) the transcriptomic matrix only, and (iii) the combined phenotypic and transcriptomic matrix. This resulted in gene pairs with corresponding correlation values for each dataset. STRING interaction scores (predicted protein links) for gene pairs were obtained through the STRING database v12 (<u>protein.links.v12.0.txt.gz).</u>

Out of the 74 genes in the CRISPRi matrix, there are 2,701 possible total gene pairs of which 1,648 had STRING interaction data available for both genes. For each dataset, the gene pairs were classified as either highly correlated (r >= 0.5) or lowly correlated (r < 0.5) and the distribution of STRING scores was visualized in box plots.

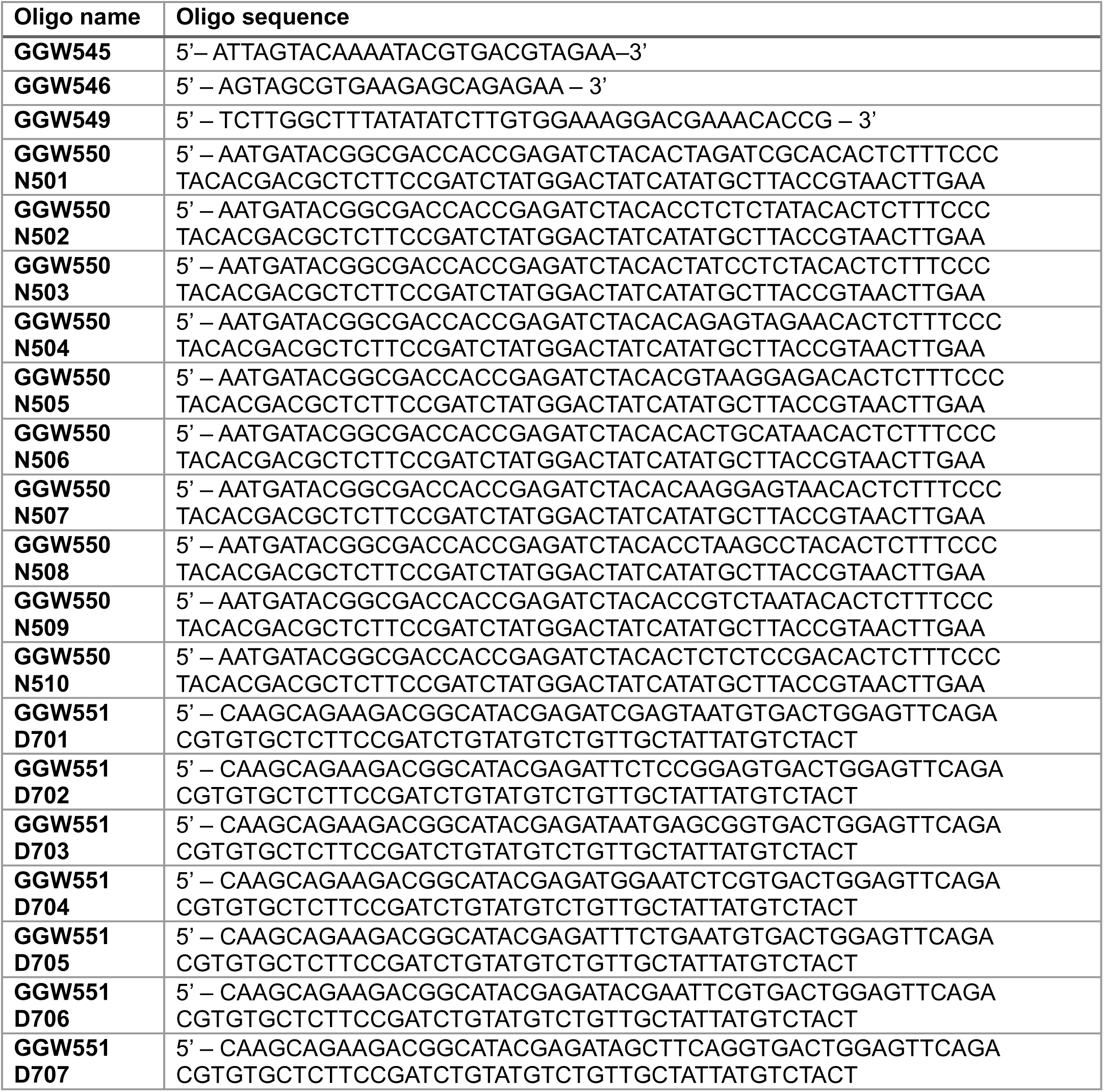
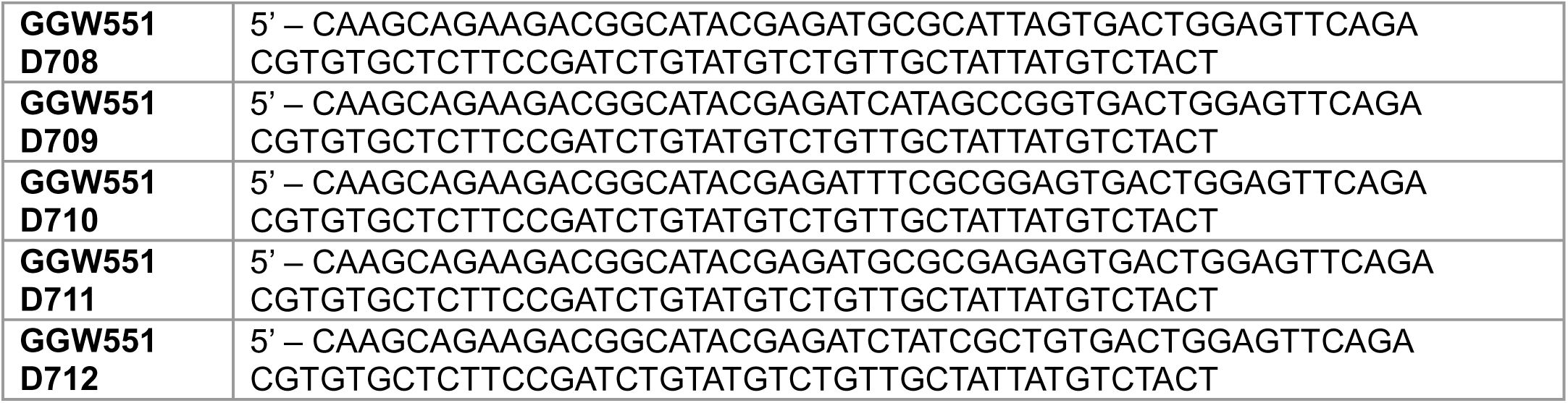

## Data and Code Availability

All code used to analyze data for this manuscript is available at https://github.com/Alector-BIO/CRISPR_PerturbSeq_pHM_publication/

Raw sequencing data and Seurat objects will be made available on GEO.

## Acknowledgements

We thank M. Kampmann, K. Srinivasan, H. Rhinn, and G. Minevich for their feedback, ideas, and suggestions on our experiments and manuscript.

## Conflict of Interest Statement

R.Y.W., E.K., H.L., S.K.M., A.R., J.A.K. are employees of Alector Inc. and may have an equity interest in Alector Inc. G.W., X.W., M.M., M.E.K., P.H., Z.K., D.R.G. were employees of Alector Inc. when contributing to the manuscript and may have equity interest in Alector Inc.

## Author Contributions

**R.Y.W.** Conceptualization, Data curation, Formal analysis, Investigation, Methodology, Software, Validation, Visualization, Writing – original draft, Writing – review and editing. **G.W.** Conceptualization, Formal analysis, Investigation, Methodology, Project administration, Resources, Supervision, Validation, Writing – review and editing. **X.W.** Investigation, Validation, Writing – review and editing. **M.M.** Conceptualization, Data curation, Formal analysis, Methodology, Software, Visualization, Writing – review and editing. **M.E.** Investigation, Methodology, Writing – review and editing. **E.K.** Investigation, Validation, Writing – review and editing. **P.H.** Conceptualization, Supervision, Writing – review and editing. **H.L.** Supervision, Writing – review and editing. **S.K.M.** Supervision, Funding acquisition, Writing – review and editing. **A.R**. Supervision, Funding acquisition, Writing – review and editing. **Z.K.** Conceptualization, Project administration, Supervision, Writing – review and editing. **J.A.K.** Conceptualization, Project administration, Supervision, Writing – review and editing. **D.R.G.** Conceptualization, Formal analysis, Methodology, Project administration, Resources, Supervision, Validation, Visualization, Writing – original draft, Writing – review and editing.

## Acknowledgments

We would like to thank Hervé Rhinn and Gregory Minevich for contributing early ideas to this research endeavor and helping design the sgRNA library. We would like to thank Karpagam Srinivasan and Martin Kampmann for their thoughtful and inspiring discussion throughout the project.

---

**Figure S1:**
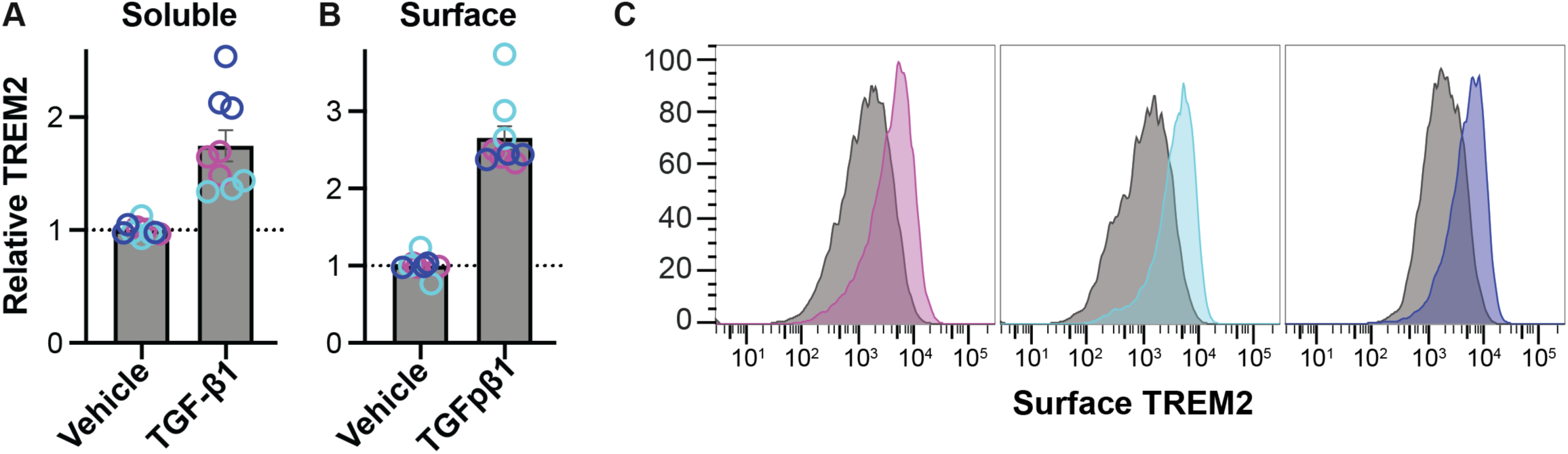
TGF-β signaling increases surface and soluble TREM2 levels. A–C) THP-1 cells were treated with vehicle or 20 ng/mL TGF-β1 for 48 h and then supernatant was collected and soluble TREM2 was measured using MSD (A) or surface TREM2 was measured by flow cytometry (B, C).

**Figure S2:**
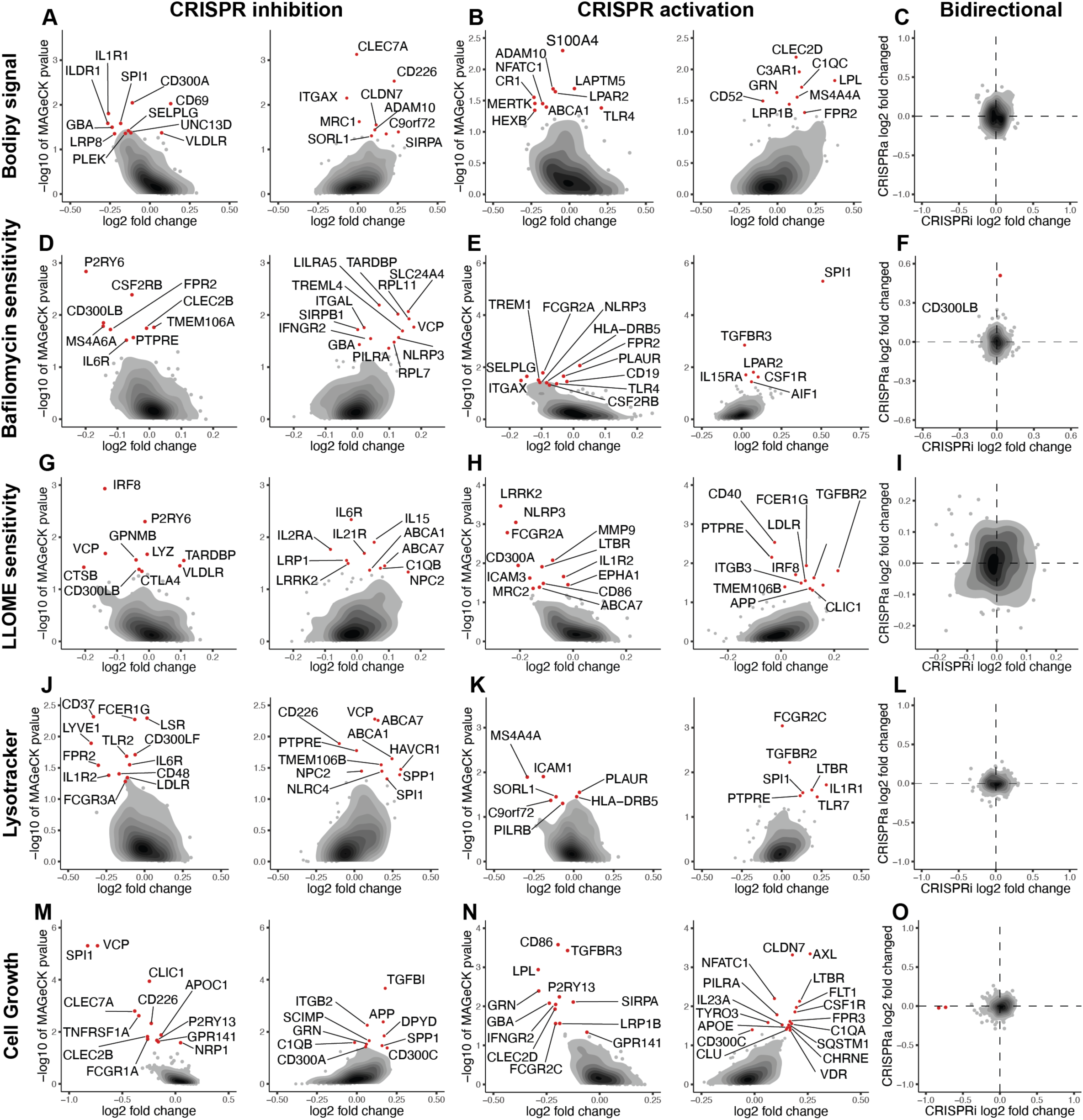
Paired CRISPRa/i screens for lipid or lysosome levels or survival in the presence or absence of lysosome stress. A, B, D, E, G, H, J, K, M, N) Dot plot with an overlayed density plot of genes from CRISPRi (A, D, G, J, M) or CRISPRa (B, E, H, K, N) screens for bodipy signal (A, B), bafilomycin sensitivity (D, E), LLOME sensitivity (G, H), lysotracker signal (J, K), or cell growth (M, N). For the fluorescent based screens (A, B, J, K) the normalized log2 fold change in readcount of the sgRNA for a given gene with the median change in abundance from the high fluorescent population relative to the low fluorescent population is on the x-axis. In the survival/growth-based screens (D, E, G, H, M, N) the change in abundance of the median sgRNA is shown after treatment relative to growth without a challenge (D, E, G, H) or relative to the sgRNA population before expansion (M, N). In both fluorescent and survival/growth-based screens the -log10 P-value calculated by MAGeCK for negative enrichment (left) or positive enrichment (right) is on the y-axis using the same comparison as defined for the x-axis. Genes with a P-value >0.05 are labeled red. C, F, I, L, O) Log2 fold change as defined above for CRISPRi screens (x-axis) plotted against the log2 fold change in the CRISPRa screens (y-axis).

**Figure S3:**
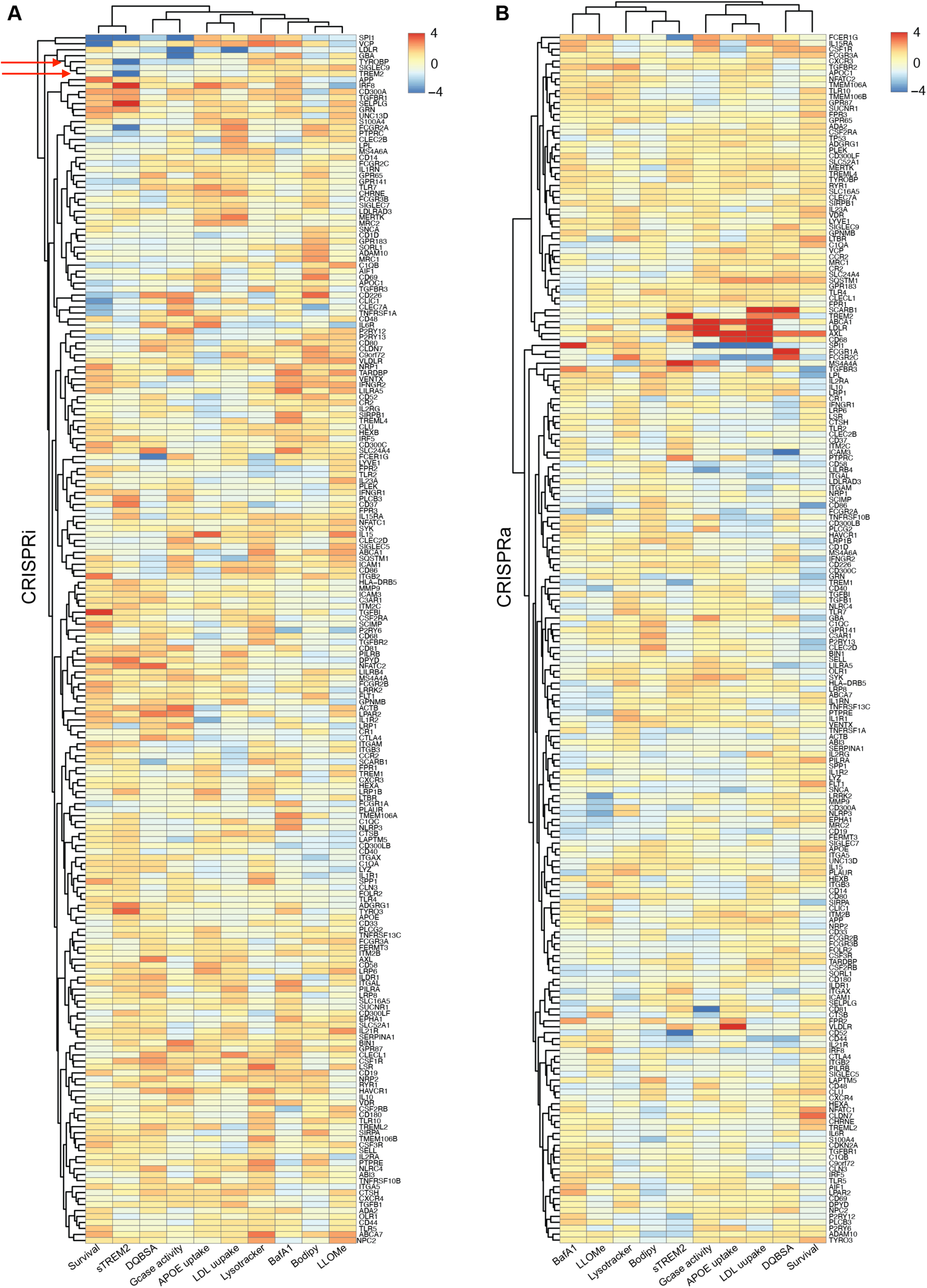
Hierarchical clustering of functional assay data in THP-1 cells. A and B) Heatmaps displaying Z-scores calculated from MAGeCK-derived p-values and log2 fold changes for CRISPRi (A) and CRISPRa (B) perturbations. Hierarchical clustering was applied to both rows (perturbations) and columns (functional assays) of the heatmaps to group similar profiles across both dimensions. Red arrows in (A) highlight the similarity of functional signatures of the co-receptors, TREM2 and TYROBP, following CRISPRi perturbation.

**Figure S4:**
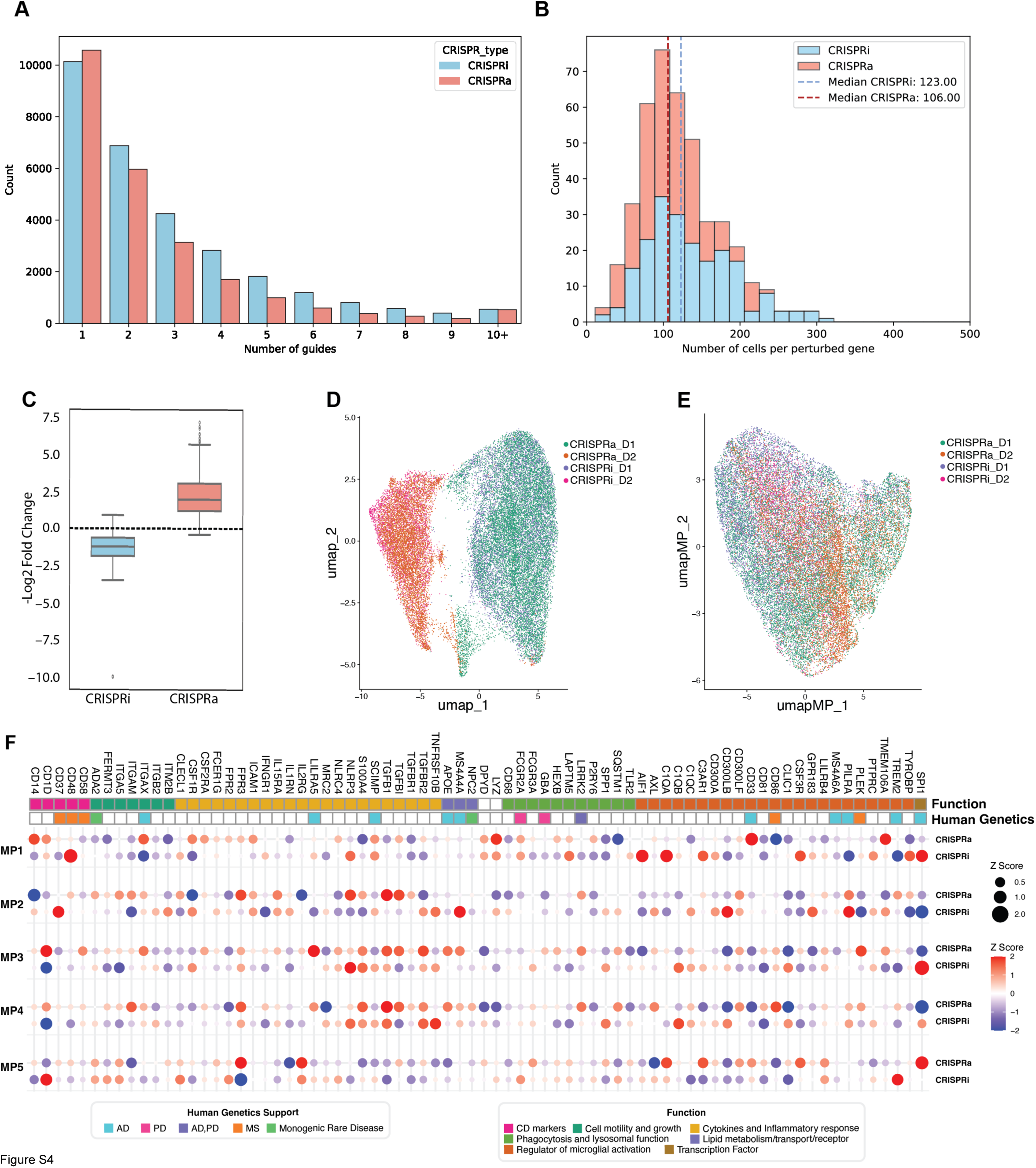
Additional QC of Perturb-seq data in pHMs. A) Bar plot showing the number of guides assigned per cell for CRISPRa (pink) and CRISPRi (blue). B) Histogram showing the number of cells per perturbed gene (inclusive of all 5 guides/gene). The distribution is shown for both CRISPRa (pink) and CRISPRi (blue), with dotted lines denoting respective median values for each library. C) Box plot showing the log2 fold changes for target genes in the CRISPRa (pink) and CRISPRi (blue) libraries. D and E) UMAP plots of CRISPRa/i libraries without donor effect correction (D) and after donor effect correction showing an integrated dataset (E). The donor and CRISPR modality were included as covariates for an integrated dataset to be used for identification of MPs through non-negative matrix factorization (NMF). F) Heatmap showing Z-transformed module scores for each MP for CRISPRa and CRISPRi perturbations. A larger circle in the heatmap indicates a stronger effect of the perturbation on the module, with the color representing the sign of the effect (red for activation and blue for repression). Perturbations are annotated by their function and human genetics support (shown in Figure 1A).

**Figure S5:**
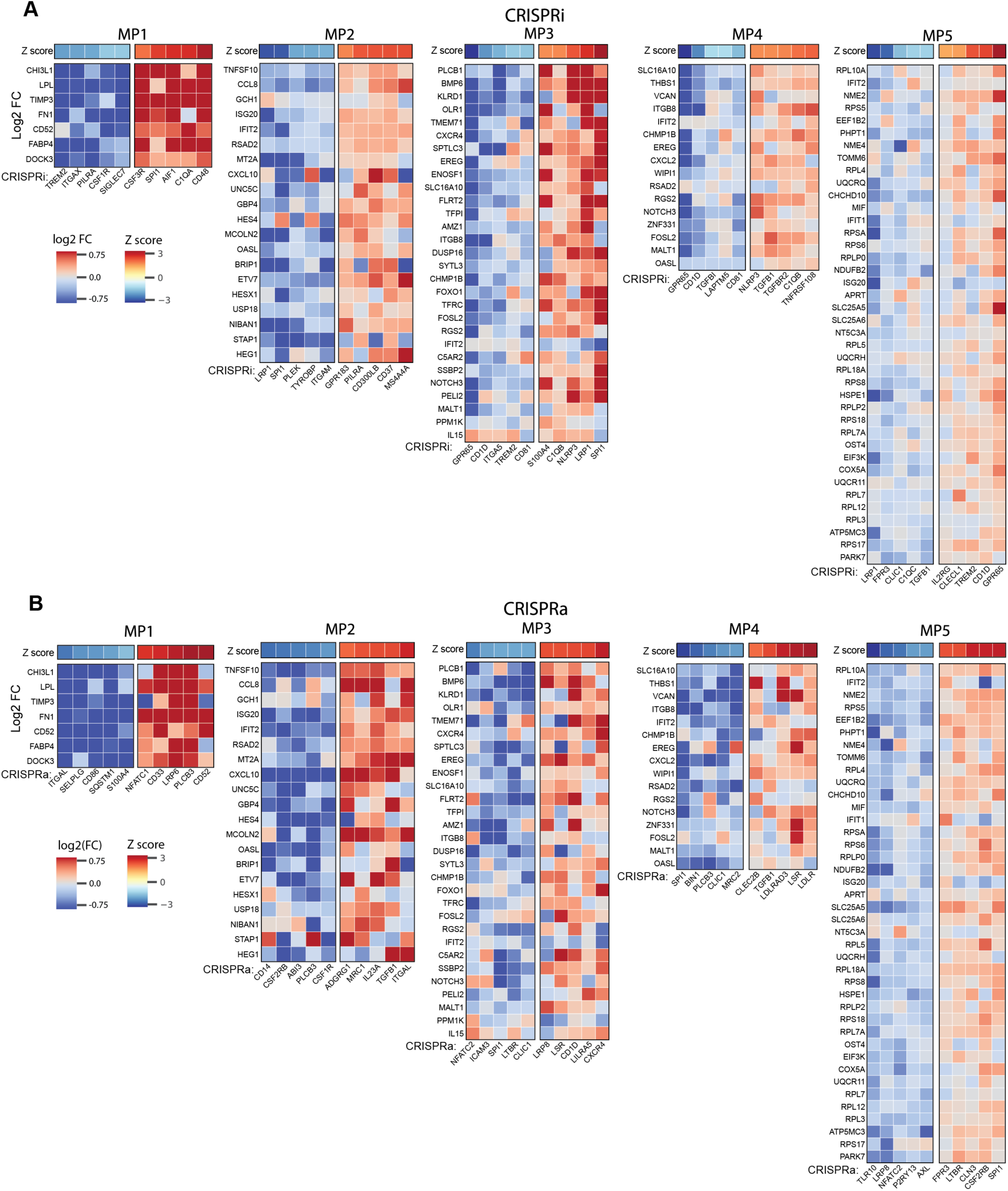
Summarized transcriptional effects on metaprograms. A and B) Extension of heatmaps in Figure 3F including all genes within each metaprogram. Colors denote log2 fold changes compared to NT control of top 5 activating and repressing perturbations in CRISPRi (A) and CRISPRa (B) libraries. Top row represents Z transformed modules scores for each MP.

**Figure S6:**
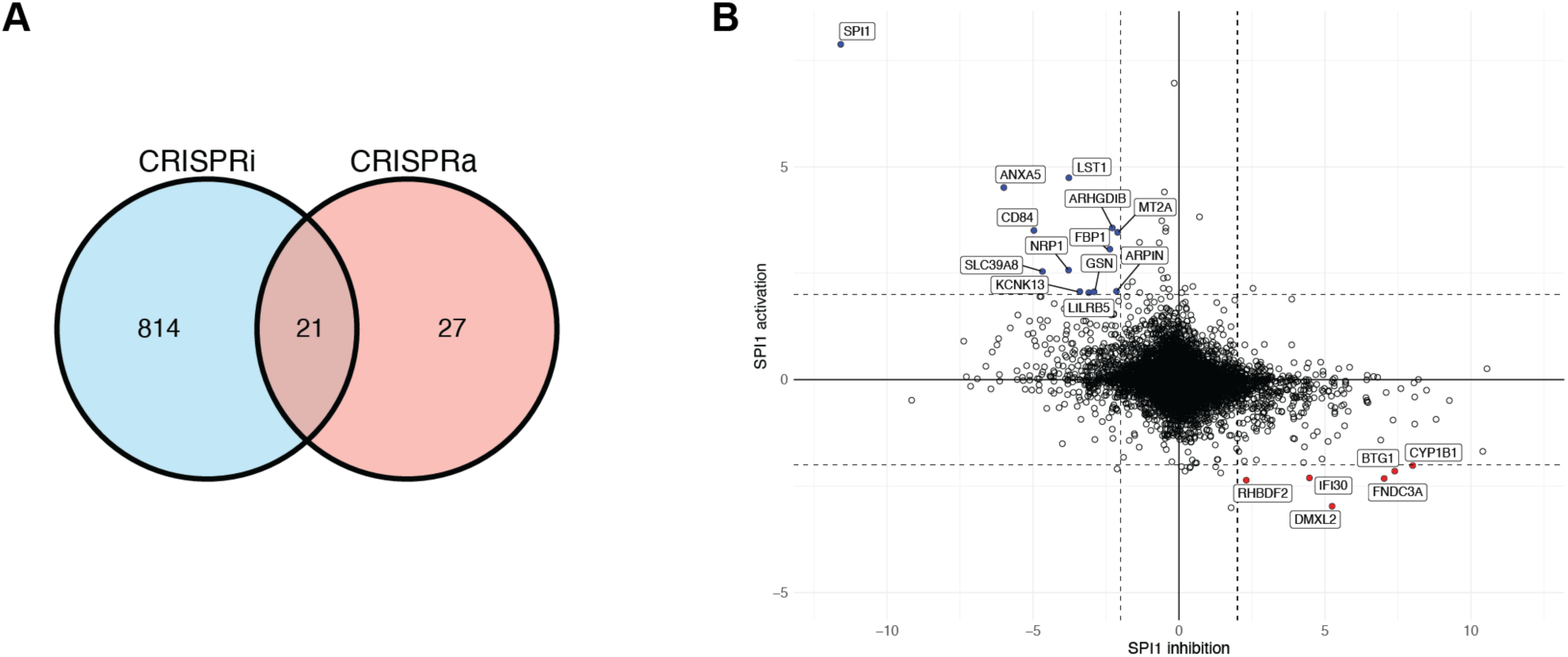
Additional effects of SPI1 perturbations. A) Venn diagram showing shared significant (adj p-val < 0.05) DEGs between CRISPRa and CRISPRi of SPI1. B) Four-way plot showing DEGs with SPI1 inhibition (x-axis) vs. SPI1 activation (y-axis).

## Notes

### Competing Interest Statement

R.Y.W., E.K., H.L., S.K.M., A.R., JAK are employees of Alector, LLC and may have an equity interest in Alector, LLC. G.W., X.W., M.M., M.E.K., P.H., Z.K., D.R.G. were employees of Alector, LLC when contributing to the manuscript and may have equity interest in Alector, LLC.

